# Improved Accuracy for Modeling PROTAC-Mediated Ternary Complex Formation and Targeted Protein Degradation via New *In Silico* Methodologies

**DOI:** 10.1101/2020.07.10.197186

**Authors:** Michael L. Drummond, Andrew Henry, Huifang Li, Christopher I. Williams

## Abstract

Extending upon our previous publication (Drummond and Williams, *J. Chem. Inf. Model*. **2019**, *59*, 1634), in this work two additional computational methods are presented to model PROTAC-mediated ternary complex structures, which are then used to predict the efficacy of any accompanying protein degradation. Method 4B, an extension to one of our previous approaches, incorporates a clustering procedure uniquely suited for considering ternary complexes. Method 4B yields the highest proportion to date of crystal-like poses in modeled ternary complex ensembles, nearing 100% in two cases and always giving a hit rate of at least 10%. Techniques to further improve this performance for particularly troublesome cases are suggested and validated. This demonstrated ability to reliably reproduce known crystallographic ternary complex structures is further established through modeling of a newly released crystal structure. Moreover, for the far more common scenario where the structure of the ternary complex intermediate is unknown, the methods detailed in this work nonetheless consistently yield results that reliably follow experimental protein degradation trends, as established through seven retrospective case studies. These various case studies cover challenging yet common modeling situations, such as when the precise orientation of the PROTAC binding moiety in one (or both) of the protein pockets has not been experimentally established. Successful results are presented for one PROTAC targeting many proteins, for different possible PROTACs targeting the same protein, and even for degradation effected by an E3 ligase that has not been structurally characterized in a ternary complex. Overall, the computational modeling approaches detailed in this work should greatly facilitate PROTAC screening and design efforts, so that the many advantages of a PROTAC-based degradation approach can be effectively utilized both rapidly and at reduced cost.

## INTRODUCTION

PROTACs (proteolysis-targeting chimeras) are heterobifunctional small molecules comprised of two binders connected by a (generally flexible) linker. The simultaneous binding of a PROTAC to both a target protein and an E3 ligase facilitates ubiquitination, the naturally occurring process whereby proteins are not merely *inhibited*, but instead are *degraded* into their constituent amino acids.^1^ In the ten years since the first small molecule PROTAC was reported,^2^ this modality of targeted protein degradation (TPD) has evolved from a chemical biology tool capable of enabling chemical protein knockdown to an approach with bona fide drug discovery applications – culminating (thus far) with Arvinas’ ARV-110 and ARV-471 entering into clinical trials for prostate and breast cancer, respectively.^3^ As a result of a sustained research effort across academia and industry, over 50 unique proteins have been degraded to date using PROTACs,^4–6^ including targets in areas of intense pharmaceutical interest such as viral proteins^7^ and aberrant tau.^8^

Interest in PROTAC-mediated TPD is driven by a desire to exploit the inherent advantages of a degradation-based modality. Perhaps the most tantalizing of these advantages is that the target-binding moiety of a PROTAC need not possess exquisite potency against its target, as it must bind only just enough for the desired effect (*i*.*e*., the degradation of the target) to occur. This lessened potency requirement vis-à-vis a more traditional protein inhibition approach is crucial for successfully disrupting *e*.*g*., scaffolding proteins^9,10^ or protein-protein interactions,^11^ where weak binders are often all that can be developed. Additionally, because PROTAC-mediated degradation occurs via the formation of a ternary, or three-body complex (PROTAC, target protein, and E3 ligase), enhanced selectivity typically results as a consequence of this additional machinery. Indeed, this phenomenon is so commonplace that Jiang *et al*.^12^ pursued a PROTAC approach specifically to selectively target either cyclin-dependent kinase (CDK) 4 or CDK6, necessitated by the tight conservation of the residues in the kinase ATP binding site. The duration of a PROTAC-mediated response has also been shown^13^ to be both immediate and long-lasting, due to the fact that the target is not merely inhibited, but actually destroyed, and therefore must be resynthesized to once more exhibit its deleterious effect. Finally, the catalytic nature of PROTACs, where a single molecule can signal for multiple copies of a target protein to be degraded,^14^ has obvious benefits for dosing and concomitant mitigation of potential off-target effects.

However, despite undeniable progress in TPD research, any degradation-based modality involves novel challenges – challenges above and beyond the already nontrivial task of developing *any* new drug – that must be surmounted. Recent studies^15–17^ have reported the evolution of drug resistance after repeated exposure of cancer cells to PROTAC-based treatments. Fundamentally, this resistance seems to involve mutations not to the targeted protein itself – as might be expected for an inhibition-driven approach^18^ – but rather to various components of the E3 ligase that ubiquitinates the target. Moreover, these mutations can occur in multiple components of the E3 ligase assembly: *e*.*g*. on the substrate recognition subunit for cereblon-based PROTACs (*i*.*e*. to cereblon itself), but on the cullin2 scaffold for von Hippel-Lindau (VHL)-based PROTACs. This mutability of the E3 ligase machinery seems therefore to potentially require the undesirable coadministration of multiple PROTACs utilizing different E3 ligases, or ideally the identification of an E3 ligase whose endogenous functionality is so integral to the normal operations of a cell that it is therefore resistant to evolutionary selective pressure. Towards this latter point, Schapira *et al*.^19^ have meticulously established a roadmap for expanding the known E3 ligase repertoire, which will also prove advantageous in the pursuit of tissue-specific PROTACs.^20,21^

Another fundamental challenge is the long-recognized fact that, while PROTACs are nominally small molecules, they are nonetheless *big* small molecules, occupying beyond rule-of-five chemical space.^22^ Maple *et al*.^23^ recently catalogued a number of key physicochemical properties (molecular weight, clogP, rotatable bond count, etc.) of published PROTACs, showing how the choice of both the E3 ligase binding moiety and the linker^24^ can have a tremendous impact on the likelihood for cell permeability and oral bioavailability, in addition to the obvious variability in the properties imparted by the target binding moieties. Properties of this sort have already been used as iterative design criteria,^25,26^ and obviously such considerations will be critical in establishing the pharmacokinetic/pharmacodynamic (PK/PD) relationships essential for successful development of *in vivo* studies^27^ and resulting treatments.

Fortunately, new techniques have continually been developed to address the new and emerging challenges of PROTAC development. Chemoproteomic profiling, particularly where electrophilic fragments react with cysteine, has repeatedly been used to elucidate novel E3 ligases, often leading to the rapid design of PROTACs that covalently bind to their ligases.^28–30^ (However, it has recently been suggested that such covalent binders are perhaps ill-suited for PROTACs,^31,32^ although this finding has itself been questioned.^33^) Multiple independent research groups have recently developed photoactivated PROTACs,^34–37^ and novel experimental diagnostic techniques^38–40^ will continue to afford additional insight with both greater accuracy and throughput.

In 2019 we reported^41^ a suite of computational modeling tools to facilitate PROTAC design, thus enabling the cost- and time-saving advantages of an iterative design campaign guided by *in silico* modeling results. The goal of our original work was to develop a *generally applicable* PROTAC modeling protocol, *i*.*e*., one that could give accurate results across different protein targets, E3 ligases, and linkers; such an approach had been absent from the literature at that time, with all previous PROTAC modeling studies limited to a specific TPD system (although preprints following our model have recently appeared^42,43^). To that end, we established “Method 4” as the most accurate approach for both VHL- and cereblon-based PROTACs. In Method 4, putative ternary complexes are constructed by first performing protein-protein docking of the target+binder complex against the E3 ligase+binder complex. Separately, a conformational ensemble is generated for the user-provided PROTAC, after which these two ensembles are combined and scored. The accuracy of Method 4 was first established through validation against known ternary complex crystal structures, establishing that it could be applied to the more realistic scenario where such crystal structures do not exist, and thus predictions must be judged solely by comparison to experimental measurements of protein degradation efficacy.

In this current work, we extend upon our initial publication^41^ in a number of important ways. First, a subtle but important change is implemented in Method 4 to produce Method 4B, which yields noticeably superior predictions. The accuracy of these predictions is further improved by *clustering* the results, which in some cases yields a final ensemble of predicted ternary complexes that almost entirely resembles the known ternary complex crystal structure. Additionally, a new method (Method 5) is detailed herein, representing a faster (albeit less accurate) alternative to Method 4B. Using these two new methods, we then extend the test set significantly beyond that used in our 2019 work. The seven case studies detailed below explore a number of common PROTAC modeling scenarios – for example, how to model successfully when there is no crystal structure available for the target+binder complex. We expect that not only will the specific results and techniques presented below prove useful for researchers using our proposed Methods, but moreover many of the findings discussed herein should be general and transferrable, so that the accuracy of independently developed PROTAC modeling procedures, such as those recently described by Li *et al*^44^ or elsewhere in preprint manuscripts,.^42,43^ can be further refined.

## MATERIALS AND METHODS

The two methods described in this publication, like our previously developed methods,^41^ were written in MOE’s^45^ native Scientific Vector Language (SVL) and are freely available upon request. The output generated by all of our *in silico* modeling methods is an ensemble of ternary complexes, where select conformations of user-provided PROTACs were judged (through varying means) to be able to successfully bridge between the binding pockets of the specified E3 ligase and the target protein. Successful formation of these ternary complexes is a necessary (but not sufficient) step before ubiquitin can be transferred from the E3 ligase machinery to solvent-exposed lysines on the target protein.^46,47^ Although it seems that the ternary complex need not be particularly stable or long-lived for target degradation to occur,^48,49^ it is becoming increasingly apparent that “cooperativity” – where formation of the three-body complex is favored over formation of either binary complex – is beneficial for effective protein degradation.^50,51^ It is thus expected – and indeed is borne out by the validation data (see below) – that the methods of this work should prove especially successful at modeling these more stable ternary complexes.

### Development of Method 4B from Method 4

We begin the discussion of our improved PROTAC modeling techniques by considering Method 4, which was previously shown^41^ to be successful at recapitulating a benchmark set of ternary complexes whose geometries had already been established via X-ray crystallography. For Method 4, two binary protein-ligand complexes are required: one containing an E3 ligase with its binder appropriately placed within the binding site, and a second containing the target protein to be degraded, also with its binder correctly located in its binding site. It is essential that the two binders in these complexes exactly match the binding moieties on either end of the bifunctional PROTAC(s) under consideration. The topic of how to place the two binding moieties into their respective protein pockets will be explored more fully below via case study; target+binder and ligase+binder X-ray crystal structures are ideal, as these will establish the respective binding geometries via experiment, but as will be shown these are not strictly required.

Once the two binary complexes are provided, protein-protein docking is used to generate an ensemble of how these two complexes might interact, absent the explicit linker of the PROTAC. These poses can be automatically generated on-the-fly using MOE’s^45^ own protein-protein docking algorithm. It is also possible to use a pregenerated ensemble of protein-protein docked poses, such as might be produced with an earlier run of Method 4, or via a standalone application of MOE’s protein-protein docker, or as imported from a different modeling program. Furthermore, it is also possible to automatically increase the diversity of the ensemble of protein-protein docked poses by repeated, separate runs of the docking procedure, where hydrophobic patches near the ligand pockets on each protein are matched in a pairwise manner. This option – referred to previously^41^ and below as a Biased simulation (*i*.*e*., protein-protein docking is “biased” to match exposed hydrophobicity against exposed hydrophobicity) was further extended in this present work: in addition to matching hydrophobic patches across the two proteins, a separate simulation is now also automatically performed, where the residues near the two ligand binding pockets are used to define the site of engagement without consideration of hydrophobic patches. In other words, this additional ensemble is the same generated if the Biased protein-protein docking option were *not* requested; the poses of this Unbiased simulation are automatically folded into the results of the larger Biased ensemble and are subjected to the same duplicate removal procedure previously discussed.^41^

Following protein-protein docking, the next step in Method 4 is the generation of a robust conformational ensemble for each user-provided PROTAC. It is in this step that Method 4B differs from its parent Method 4. In the original Method 4, no restrictions were placed on the PROTAC during this conformational search procedure. By contrast, in Method 4B of this work, each binding moiety in the PROTAC is constrained to retain its bound conformation, as provided in the two protein-ligand complexes (ligase+binder and target+binder) utilized in the previous protein-protein docking step. While at first glance this change may seem minor, the ramifications of this modification are many and important. Most immediately, far fewer PROTAC conformations are generated. For example, Figure 1 shows conformational ensembles generated for PROTAC 2 of Farnaby et al.^52^ using Methods 4 and 4B; the former (Figure 1b) produced 4374 conformations, whereas Method 4B generated 60 conformations (Figure 1c) using the same conformational search settings (LowModeMD^53^ allowing a maximum of 10,000 unique conformations and halting after 100 consecutive duplicate conformations). Additionally, because Method 4B produces far fewer conformations than Method 4, the final step common to both methods – the combination of the protein-protein docked ensemble and the PROTAC conformational ensemble – takes proportionally less time in Method 4B than in Method 4. Moreover, Method 4B no longer involves a subjective definition of what constitutes a “core” (as was required in Method 4), as each rigid binding moiety is defined as the respective core. Finally, the geometries of the two PROTAC binding moieties initially fit perfectly into their respective binding pockets, although some distortion of these binding geometries typically occurs during minimization of the final overlaid PROTAC conformation (*i*.*e*., the transition from e) to f) in Figure 2 of Ref. 41). Attempting to maintain the conformational restraints on the two binding moieties during this final minimization step (*i*.*e*., throughout the entire modeling process) yielded poorer overall results for the validation set of known ternary complex crystal structures (data not shown), as perhaps might be expected if the linker, as is known,^54^ can interact with the two PROTAC binding moieties and their respective pockets.

**Figure 1.**
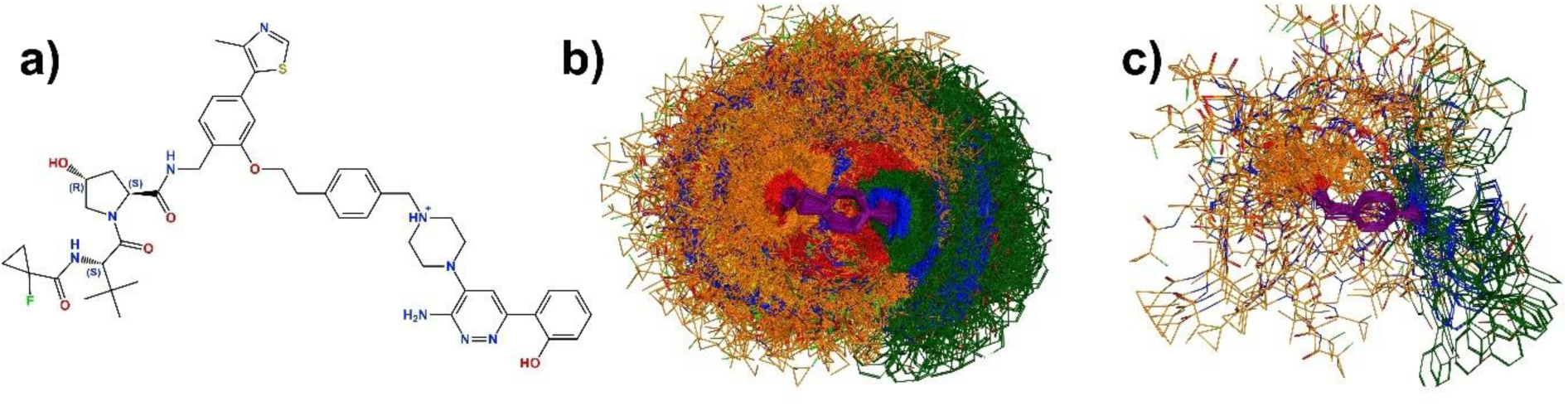
Differences in the conformational ensembles generated for a) PROTAC-2^52^ using b) Method 4 and c) Method 4B. In b) and c), conformations were superposed on the *p*-ethylmethylbenzene linker (purple) of PROTAC-2, with the carbons of the VHL and SMARCA2 binding moieties colored orange and green, respectively. Visual clipping was used in b) to provide a glimpse of the purple linker core; absent such clipping, there is a complete “shell” of orange and green atoms surrounding the linker. No such clipping was required for c).

**Figure 2.**
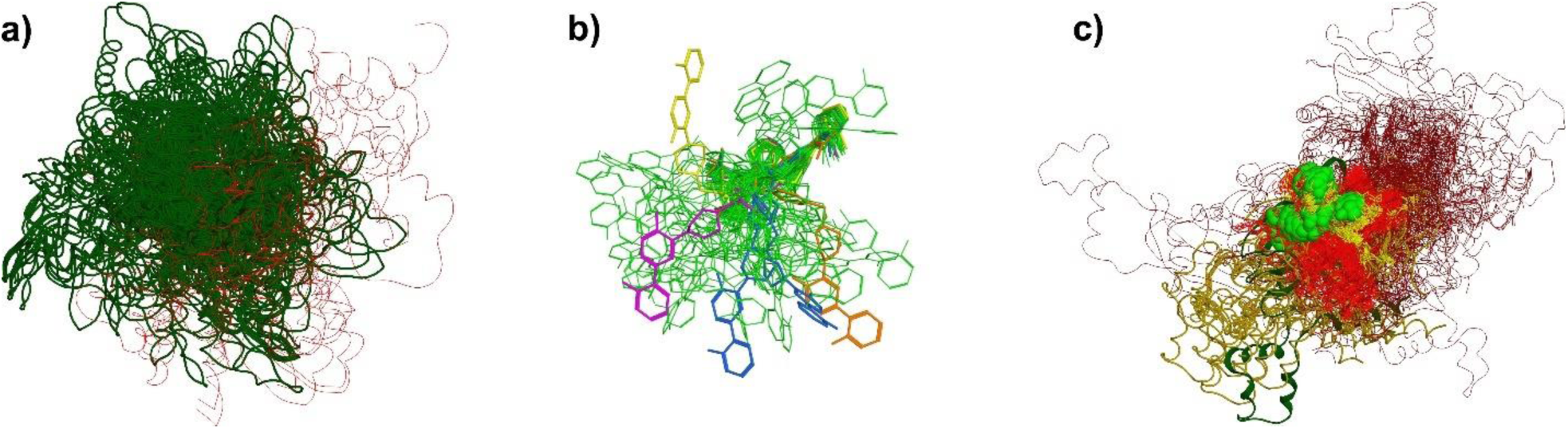
a) Modeled ternary complexes produced with Method 4B for 6HAX,^52^ shown top-down from the perspective of the E3 ligase VHL (which was used as a common frame of reference for superposition). For clarity, only the ribbons of SMARCA2^BD^ are shown. The dark green ribbons belong to the largest double cluster (see text), whereas those in red belong to other clusters. None of the red ribbons shown are within 10Å of the crystallographic positions of SMARCA2^BD^ in 6HAX. b) Conformations of PROTAC-2^52^ produced with Method 4B that form ternary complexes (after filters are applied) for 6HR2. These conformations have been superposed on the VHL binding moiety (thick grouping in the upper-right corner) and have been colored by their cluster identity. The green conformations all belong to the same cluster, which is the most populous cluster for PROTAC-2. c) Modeled ternary complexes produced with Method 4B for 6HR2, shown top-down from the perspective of the target protein SMARCA4^BD^ (used as a common reference for superposition and hidden for clarity). The ribbons shown are for various poses of the E3 ligase VHL. The dark green ribbon – actually 49 copies of the same protein-protein docked pose (see Table S1) – represent the position of VHL in the complexes belonging to the largest double cluster, all of which superpose to within 10Å of 6HR2. The golden ribbons (123 proteins) also belong to the largest double cluster, but do not superpose within 10Å of 6HR2. The dark red ribbons do not belong to the largest double cluster, nor do they superpose within 10Å of 6HR2. The conformations of PROTAC-2 are also shown, in light green (and space filling), yellow, and bright red, corresponding, respectively, to the dark green, golden, and dark red ribbons, respectively.

### Validation of Method 4B

Having established the methodological details of Method 4B, we now turn to validating its performance, especially through comparison to the accuracy already established^41^ for Method 4. As before, this initial validation involves judging the ability to reproduce ternary complex geometries as revealed through structures elucidated by X-ray crystallography. The six crystal structures used for this validation in this current work are listed in Table 1. Note that the structures in PDB codes 6BN8 and 6BN9,^55^ which were included in our original validation set,^41^ were removed from consideration in this study, as their low resolutions precluded the identification of any part of the PROTAC (and as new ternary complex crystal structures have since been published,^52^ and thus could be swapped in). A seventh ternary complex crystal structure, 6SIS,^56^ was only recently published and therefore was not included in the validation set, although results for modeling this crystal structure will be discussed below as Case Study 1. As part of this validation effort, new “filters” – *i*.*e*., properties calculated on the modeled ternary complexes with accompanying thresholds – were evaluated, based on their ability to distinguish modeled ternary complexes that satisfactorily “resemble” the X-ray crystal structures from those that do not. Two criteria were used to judge this resemblance to the crystal structures (Table 1): first, as before,^41^ whether or not the modeled ternary complex could be superposed as a rigid body to the corresponding crystal structure with a total C_α_ RMSD of ≤10Å; and second, using the high/medium/acceptable criteria of the CAPRI^57^ protein-protein docking assessment, which can consider a pose to be “acceptable” if the interfacial residues are largely maintained, even if the distal end of one of the proteins does not superpose well, and therefore the global C_α_ RMSD exceeds 10Å.

**Table 1.** Validation Results for Methods 4 and 4B for Known Ternary Complex Crystal Structures.

| PDB | E3 Ligase | Method 4 | Method 4B |  |
| --- | --- | --- | --- | --- |
|  |  | Crystal-Like/Total (Hit Rate) | Crystal-Like/Total (Hit Rate) | High/Medium/Acceptable (Hit Rate)<sup>a</sup> |
| 5T35<sup>47</sup> | VHL | 68/172 (39.5%) | 979/1692 (57.9%) | 50/950/198 (70.8%) |
| 6BN7<sup>55</sup> | Cereblon | 2/23 (8.7%) | 15/390 (3.8%) | 0/5/15 (5.1%) |
| 6BOY<sup>55</sup> | Cereblon | 0/13 (0.0%) | 4/1063 (0.4%) | 0/4/0 (0.4%) |
| 6HAX<sup>52</sup> | VHL | 0/591 (0.0%) | 143/526 (27.2%) | 88/36/71 (37.1%) |
| 6HAY<sup>52</sup> | VHL | 309/805 (38.4%) | 79/420 (18.8%) | 0/36/144 (42.9%) |
| 6HR2<sup>52</sup> | VHL | 49/534 (9.2%) | 65/447 (14.5%) | 0/67/32 (22.1%) |

<sup>a</sup>Across all three categories

The new filters for Method 4B were selected from the set of 70+ properties detailed in the Supporting Information of Ref. 41, including metrics quantifying how much of one protein’s surface patch area (be it hydrophobic, negative, and/or positive) is covered by similar protein patches on the second protein as the ternary complex comes together. In this current study, for Method 4B it was decided to move away from multiple sets of filters – *cf*. Method 4^A^ – 4^F^ in Table 1 of Ref. 41 – in favor of a more widely applicable “one-size-fits-all” approach. Thus, the results in Table 1 of this publication were generated using Method 4B with the same two filters always applied. The first such filter, Core RMSD ≤ 3.5 Å, measures the goodness-of-fit between each PROTAC conformation and each protein-protein docked pose, evaluated exactly as described previously.^41^ The larger value of 3.5 Å (*cf*. a previous maximum of 1.6 Å for Method 4^F^)^41^ is a consequence of the expansion of the definition of the eponymous “core” to now encompass the entirety of the target and E3 ligase binders. The second filter used for the results in Table 1 describes the *total* direct protein-protein interfacial surface area – *i*.*e*., not limited to interfacial protein *patches* – of the two proteins in a predicted ternary complex. Specifically, only ternary complexes with a total interfacial surface area of *less than* 670 Å^2^ are considered, with all other putative ternary complexes discarded. *A priori*, this threshold is likely a consequence of the fact that the target protein in all six ternary complex crystal structures is a bromodomain (BD), of either Brd4, SMARCA2, or SMARCA4. These BDs are anisotropic, binding to their corresponding E3 ligases via one “end” of a roughly cylindrical bundle of alpha helices (see Figure S1 in Supporting Information). Thus, it seems that an upper limit of 670 Å^2^ of direct protein-protein interface merely serves to select for this type of end-on binding arrangement, while poses where the target protein engages the E3 ligase in a more side-on fashion are rejected. Similarly, the two E3 ligases found across the six known ternary complex crystal structures of Table 1, *viz*. VHL and cereblon, also seem to bind their target proteins via a narrower “end,” as would also be enforced by this threshold (see Figure S1). Nonetheless, this empirically derived characterization of direct protein-protein interactions should be revisited, and revised if necessary, once additional, more varied crystal structures are made publicly available. In the meantime, we will demonstrate that useful modeling results can still be generated for non-BD targets and even for systems utilizing a different E3 ligase (cIAP1),^58^ suggesting (although by no means ensuring) that the “end-on” binding dictated by this threshold may be a general feature of PROTAC-mediated ternary complexes.

Turning now to the results in Table 1, it can be seen that Method 4B generally – although not universally – improves upon the quality of the modeled ternary complexes produced with Method 4. Four of the six crystal structures are better reproduced, as measured by the crystal-like hit rate, with Method 4B than with Method 4, with 6BN7 and 6HAY being the exceptions. However, it should be noted that Method 4 failed to generate *any* ternary complex models that superposed within 10Å to the known crystal structure for two systems, 6BOY and 6HAX. Conversely, Method 4B provided at least some correct poses in the final output ensemble for all six known crystal structures. Using the CAPRI-like high/medium/acceptable scoring criteria, where a correct definition of the interfacial residues can “rescue” a pose that would have otherwise been rejected based on a globally poor superposition, improves (or at least does not worsen [6BOY]) the hit rate, sometimes significantly (*e*.*g*. 18.8% to 42.9% for 6HAY). Arguably, correct modeling of the interface is more critical for iterative structure-based PROTAC design than a satisfactory positioning of the entire complex, as engineering improved interactions between the PROTAC and the interfacial residues is known to be a tangible design criterion.^52^

Nonetheless, it is clear from Table 1 that Method 4B yields better results for VHL-based ternary complexes than for cereblon-based complexes, as only 0.4% (6BOY) or 5.1% (6BN7) of the generated ensembles for the latter acceptably compare to the known crystal structures. However, this overall poor performance may be attributable to greater difficulty in generating initial protein-protein docking poses that resemble the final protein arrangement in the cereblon-containing ternary complex crystal structures; a recent preprint raised this same possibility (although, with their methodology, the challenging E3 ligase was VHL rather than cereblon).^42^ To test this possibility, we manually performed 63 individual protein-protein docking runs, each generating up to 100 unique protein-protein docked poses, where the seven solvent-exposed residues within 4.5Å of the cereblon binder in 6BOY were each matched up against each of the nine residues similarly near the Brd4^BD1^ binder in 6BOY. (For comparison, the automated Biased docking option described above performed 10 individual protein-protein docking runs). Without parallelization, on a typical laptop, each individual docking run takes ∼one hour, and thus this manual “Multidock” approach is likely too unwieldy for routine use. Nevertheless, this thorough Multidock approach yielded a total final ensemble of 1604 ternary complexes (*cf*. 1063 for 6BOY in Table 1), and, importantly, 229 of them (14.3%) superposed against 6BOY to within 10 Å. Therefore, it seems that the infamous “plasticity” of cereblon-based ternary complexes^55^ represents a challenge, albeit not insurmountable, for modeling efforts utilizing protein-protein docking. However, as will be illustrated by the case studies below, it seems that considering the *relative* ability of two or more cereblon-engaging PROTACs to degrade target proteins can also help ameliorate this challenge, even without the extra computational cost of this expensive Multidock procedure.

### Improving Hit Rates via Clustering

During the development of Methods 4 and 4B, it soon became apparent that the final modeled ensemble of ternary complexes often contained visually obvious groupings of ternary complexes – see for example the green poses (generated with Method in Figure 3 of Ref. 41, or Figures 2a and 2c below (generated with Method 4B). In the case of the former example, the most noticeable of these groupings also superposes well to the known ternary complex crystal structure of 5T35 (*i*.*e*., it seems enriched in crystal-like poses). To explore whether this link between crystal-like pose enrichment and visually obvious groupings of poses is general, clustering was applied to the final output ensemble of modeled ternary complexes, in hopes that perhaps the most populous cluster might generally be enriched for crystal-like poses.

**Figure 3.**
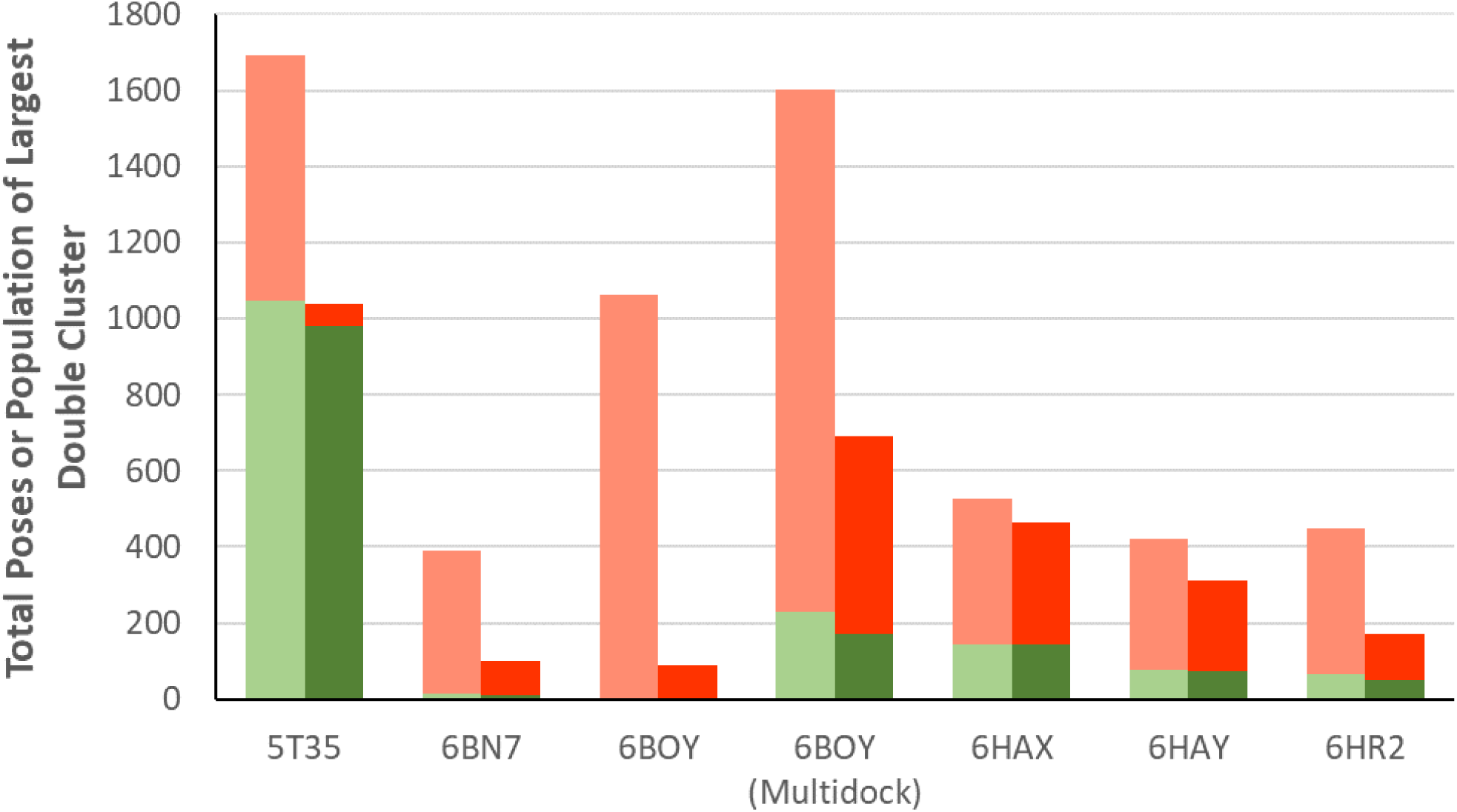
Method 4B results before and after clustering. The lighter bars correspond to the total number of unclustered poses (Table 1), whereas the accompanying darker bars show the populations within the largest L3xP10 double cluster. Poses that superpose within 10Å RMSD of the crystal structure are colored green, while those that do not are colored red. The 6BOY (Multidock) data is for the non-standard Method 4B simulation where each binding site residue in cereblon was manually paired against each binding site residue in the target protein (see text). Full numerical results are provided as Supporting Information (Table S1).

Two separate clustering algorithms were applied to the final ensemble of modeled ternary complexes: one to the proteins themselves and one to the PROTAC conformations used in these ternary complexes. For the proteins, MOE’s implementation of single-linkage/nearest-neighbor clustering was utilized, via the SVL function hclust_tree, based on the C_α_ RMSD of the moving protein after rigid-body superposition on the stationary protein. Clustering thresholds of 5, 10, and 15Å were explored. Similarly, the PROTAC conformations were assigned to clusters using a nearest-neighbor clustering algorithm based on the all-against-all conformer-to-conformer heavy atom RMSD matrix after ideal rigid-body superposition. This algorithm also accounts for graph isomorphisms as the conformers are superposed. For this clustering, thresholds of 1-3Å in 0.5Å increments were explored.

Cluster numbers were thus generated separately for the protein and PROTAC conformations found across all poses in the final Method 4B output of ternary complexes. From here, a “double cluster” number was assigned simply by concatenating the two individual cluster numbers. For example, a ternary complex with a double cluster number of 2_4 simply means that the PROTAC conformation contained in that ternary complex was clustered into ligand cluster 2, while the protein poses of that complex were assigned to protein cluster 4. (Note that there is not necessarily an ordering of these individual cluster numbers by population, *i*.*e*., cluster 1 is not necessarily the largest cluster). After generating all double cluster numbers for each output ternary complex, the population of each unique double cluster number was tabulated. This procedure was repeated across all possible combinations of the two individual cluster thresholds – *i*.*e*., 5, 10 and 15Å *x* 1, 1.5, 2, 2.5, and 3Å for the protein and PROTAC clusters, respectively. The specific thresholds for the optimal double clustering procedure were chosen based on which combination of thresholds yielded the best hit rate of crystal-like poses in the most populous double cluster, as applied to the validation dataset of Table 1. Additionally, clustering based only on the proteins or only on the PROTACs (*i*.*e*., the single clusters) was also investigated, although these did not outperform the best double clustering procedure (data not shown).

The best enrichment in the hit rate across the six structures of the validation set was found in the most populous double cluster built from a 3Å clustering threshold for the PROTAC ligands and a 10Å threshold for the proteins themselves, hereafter referred to as the L3xP10 double cluster. Figure 3 shows the impact on the hit rate (as measured by counting poses that can superpose to the known crystal structure within 10Å RMSD) of disregarding any poses that do not belong to this most populous double cluster; Table S1 in the Supporting Information instead shows the same effect considering the CAPRI-like definition of hit rate instead. As can be seen, clustering generally reduces the number of false positives as the raw output ensemble is restricted to the largest double cluster (light and red bars, Figure 3). Occasionally, some true positives are grouped in a cluster separate from the largest double cluster – for example, for 6HR2, 16 crystal-like poses do not belong to the largest double cluster (15 belong to the 4^th^-largest cluster, and one is a singleton) – and thus these desirable true positive results are discarded when only the largest double cluster is considered. While it is certainly possible to evaluate smaller double clusters as well, as has been done elsewhere,^42^ unless additional experimental information is available for use to evaluate these clusters, it seems likely that more noise than signal will generally result from this more expansive approach. Moreover, as can be seen in Figure 3, even if some crystal-like poses fall outside the largest double cluster, the consequence of successfully discarding false positive ternary complexes almost always outweighs this minimal loss of true positives, and thus this double clustering procedure improves the hit rates in nearly all cases. The sole exception is for 6BOY, where only 0.4% of the unclustered ternary complex ensemble is crystal-like (Table 1), and none of these four true positives are lumped into the largest double cluster. However, as discussed above, this poor showing results from generating an unsuitable protein-protein docked ensemble that contains few crystal-like poses in the first place. If a larger, more expansive protein-protein docked ensemble is clustered (*i*.*e*., the Multidock results of Figure 3), then 25% of the most populous double cluster superposes well with 6BOY – an improvement of over 10% in the hit rate compared to the unclustered Multidock results.

Figure 2 visually depicts some of the different ways the double clustering procedure improves the hit rate in the final ensemble of ternary complexes. In Figure 2a, depicting modeled results for 6HAX, all conformations of PROTAC-2 which successfully form a ternary complex that passes the two Method 4B filters (see above) belong to the same ligand cluster. Thus, considering only the largest double cluster in this case is simply considering only population of the largest protein-based cluster. There are 62 proteins that do not belong to this largest protein cluster (red ribbons in Figure 2a), and none of them superpose to within 10Å of 6HAX. Thus, discarding the poses that belong to a smaller double cluster merely “trims the fat” away from a core ensemble of poses that generally resembles the crystallographic positioning, albeit with some “fuzziness” (*i*.*e*. 40.1% of this trimmed-down ensemble resembles the 6HAX crystal structure based on the interface-favoring CAPRI-like assessment criteria). For 6HR2, the PROTAC conformations found in all complexes across the final ternary complex output are assigned to one of five clusters, one of which (the green conformers of Figure 2b) is much more populous than the others; the protein poses in the final ternary complexes likewise are clustered into 10 separate clusters. Importantly, 65 crystal-like ternary complexes in the final ensemble are divvied into only three protein clusters after the nearest-neighbor algorithm is applied, which contain 49, 15, and 1 crystal-like poses. The dark green ribbons in Figure 2c are the 49 poses (actually 49 copies of the same protein-protein docked pose) linked by different but similar PROTAC conformations (*i*.*e*., they all belong to the same, most populated ligand cluster). This largest double cluster also contains 123 ternary complexes whose proteins superpose to 6HR2 with an RMSD of more than 10Å (although always <18Å), which are depicted with golden ribbons in Figure 2c. Ternary complexes that do not belong to this largest double cluster and that do not superpose well with 6HR2 are shown in red, which as can be seen clearly differ from the green and gold ribbons of the largest double cluster. Thus, discarding these poses removes a great many non-crystal like false positive poses, thereby nearly doubling the hit rate for 6HR2 (from 14.5% to 28.5%).

Overall, this double clustering procedure produces a focused pool of modeled ternary complexes, substantially enriched for crystal-like poses beyond the results of any of our other modeling Methods. For those ternary complexes that use VHL as their E3 ligase, over 40% of the poses in the most populous double cluster are at least “acceptable” when compared against the known crystal structure – culminating in the performance for 5T35, where almost *every* modeled complex (99.3%) in the largest double cluster is acceptable or better by the CAPRI standards. For the two cereblon-utilizing ternary complexes, if the protein-protein docking algorithm successfully generated crystal-like poses (*i*.*e*., considering the Multidock results for 6BOY), the double clustering approach will still yield a largest double cluster with a double-digit percent population of crystal-like poses. Due to these improved success rates, all case studies considered in this work using Method 4B will be scored using the total population of the largest double cluster (L3xP10). In practice, however, to guard against a situation encountered with the Biased (but not Multidocked) 6BOY simulation, where clustering assigned the few identified crystal-like poses to a smaller double cluster, the final output given by the Method 4B script is the entire *unclustered* set of ternary complexes, with the corresponding double cluster identities and populations indicated in additional columns. This expansive output will also facilitate investigations into alternative clustering approaches.

### Method 5 – Spanning Ligand Pockets *in situ* with PROTACs

Finally, below we propose and validate another method for *in silico* modeling of ternary complexes, guided by a desire to exploit the ensemble of protein-protein docked poses generated with Methods 4 and 4B in a more efficient manner. As such, Method 5 also beings with the protein-protein docking of two protein-ligand complexes (target+binder and E3 ligase+binder). Moreover, as in these two extant methods, protein-protein docking ensembles can be generated on-the-fly with the script – both with and without the “Biasing” procedure described above and earlier^41^ – or can be reused, either from earlier simulations (of Methods 4 or 4B, or from previous runs of Method 5) or after importing from third-party protein-protein docking algorithms.

After an ensemble of protein-protein docked poses is generated or supplied, the PROTAC linker, which is not included in the protein-protein docking simulation, must be spanned between the two protein binding moieties. Whereas Methods 4 and 4B perform a detailed search of the PROTAC to identify conformations that can effectively span between the two binding pockets, Method 5 seeks to bridge this gap without this lengthy search. In order to find a path for the linker that can connect these pockets without unduly impinging upon the proteins, each protein-protein docked pose is first solvated with a periodic box of pre-equilibrated water. As part of this solvation process, water molecules that clash with the two protein-ligand complexes are automatically removed, ultimately filling the channel or cavity between the two binding sites with water. (This process is automatically carried out using the SVL Solvate function). Next, the shortest path between the two connection points of the two binding moieties is located by hopping from one water oxygen to another. The PROTAC linker is then connected to its two binding moieties and placed along this shortest path, before a three-step restrained minimization procedure, previously described,^41^ is performed. A similar approach has already been shown to be effective in designing bivalent linkers for various model systems, including a PROTAC-mediated ternary complex.^59^

Generally, linkers that are too long to bridge between the two pockets in a given protein-protein docked pose will coil upon themselves, whereas linkers that are too short must stretch their bonds to accomplish the bridging. These contrasting behaviors can thus conceptually be used to identify protein-protein poses that can suitably be bridged by a given PROTAC linker, resulting in a goodness-of-fit or compatibility metric between a PROTAC’s linker and a given E3 ligase/target protein pose. Moreover, the protein-protein docked poses themselves can be characterized based on how well they resemble proteins found in known (*i*.*e*., crystallographically characterized) ternary complexes. Thus, as in the development of our earlier modeling methods, filters and accompanying thresholds were empirically derived based on their ability to disregard false positives while keeping modeled ternary complexes that superpose well (*i*.*e*., within 10 Å RMSD) against experimentally known ternary complexes.

During this investigation, two filters describing the ability of a PROTAC to successfully bridge between the pockets of a protein-protein docked pose were developed. The first, RMSD_RF, is the RMSD of the heavy atoms between a) the PROTAC as placed by the solvation-based path finding approach described above in a final output ternary complex and b) the nearest local minimum of the PROTAC after performing an unrestrained minimization without either protein present, using the default R-Field (hence _RF) solvation model in MOE. This metric is essentially a measure of the strain (by geometric distortion rather than energy) of a putative bridging PROTAC conformation, absent any consideration of the proteins. A second filter, ligand_E, is simply the total forcefield energy of the PROTAC conformation as placed in a ternary complex, evaluated with the default MOE AMBER10:EHT forcefield. These two PROTAC-centric filters were also combined with a characterization of the direct protein-protein interaction. For this, the same threshold developed for Method 4B was applied, *viz*. a total interfacial surface area of ≤ 670Å^2^ was required

All told, four filtering criteria were developed: RMSD_RF < 2.5 Å, ligand_E < 3600 kcal/mol, and well as these two paired with the protein-protein (_PP) threshold of ≤ 670Å^2^. Table 2 provides the validation results for Method 5 as applied to the same six crystal structures used to develop and validate Method 4B. Additionally, Table 2 shows results for both “Unbiased” (_U) protein-protein docking simulations, where only a single docking run was performed, as well as for “Biased” (_B) docking, where multiple simulations were performed and collated, as described above and previously.^41^

**Table 2.** Hit Rates for Method 5.

| Protein | True Positives/False Positives (Hit Rate) |  |  |  |
| --- | --- | --- | --- | --- |
|  | RMSD_RF | RMSD_RF_PP | ligand_E | ligand_E_PP |
| 5T35_U | 1/1 (50.0%) | 1/1 (50.0%) | 3/52 (5.5%) | 3/24 (11.1%) |
| 6BN7_U | 1/7 (12.5%) | 1/0 (100.0%) | 2/42 (4.5%) | 2/6 (25.0%) |
| 6BOY_U | 1/3 (25.0%) | 0/2 (0.0%) | 1/42 (2.3%) | 0/8 (0.0%) |
| 6HAX_U | 2/2 (50.0%) | 2/2 (50.0%) | 2/8 (20.0%) | 2/7 (22.2%) |
| 6HAY_U | 1/0 (100.0%) | 1/0 (100.0%) | 1/8 (11.1%) | 1/0 (100.0%) |
| 6HR2_U | 2/4 (33.3%) | 2/6 (25.0%) | 2/14 (12.5%) | 2/8 (20.0%) |
| 5T35_B | 3/9 (25.0%) | 3/9 (25.0%) | 14/177 (7.3%) | 14/87 (13.9%) |
| 6BN7_B | 0/13 (0.0%) | 0/6 (0.0%) | 1/177 (0.6%) | 1/74 (1.3%) |
| 6BOY_B | 2/41 (4.7%) | 0/24 (0.0%) | 3/302 (1.0%) | 1/152 (0.7%) |
| 6HAX_B | 4/11 (26.7%) | 4/11 (26.7%) | 6/84 (6.7%) | 6/68 (8.1%) |
| 6HAY_B | 0/8 (0.0%) | 0/8 (0.0%) | 5/137 (3.5%) | 5/81 (5.8%) |
| 6HR2_B | 3/9 (25.0%) | 3/9 (25.0%) | 4/58 (6.5%) | 4/54 (6.9%) |

As can be seen, unlike with Methods 4 and 4B, where the repeat protein-protein docking steps in the Biased simulations generally yielded improved behavior (to the point that they were the only option employed in the development of Method 4B), the *Un*biased simulations almost always give higher hit rates for Method 5. We traced this unexpected finding to the inability of the added water molecules to trace out an allowed path for the PROTAC linker if the putative connecting channel is especially narrow. In some crystal-like poses, the water molecules cannot pack along the entire length of the channel, resulting in a gap in the placed waters. If this gap is larger than 5Å, Method 5 disregards the complex, judging the pathway to be occluded. While false positive complexes (which do not superpose well to known ternary complex crystal structures) are also rejected by this logic, some true positives are rejected in this fashion as well, and thus the additional, potentially useful protein-protein docked poses generated by the lengthier Biased simulations are often simply discarded. In an attempt to obviate this limitation, an artificial solvent system comprised of non-interacting argon atoms spaced evenly 1Å apart was designed and utilized. This system indeed was found to be more successful at locating tight channels through which PROTAC linkers could span. However, this more closely packed solvent also, unfortunately, tended to provide linker paths that impinged too closely upon the proteins. While conceptually it would be possible to sample multiple potential paths for the linker, rather than simply selecting the single shortest path, such a search would decrease the computational efficiency of Method 5, thereby neutralizing one of its key advantages. Thus, in the case studies presented below, results are presented using Unbiased protein-protein docking runs with the normal water solvation procedure, as a balance between expediency and accuracy.

As can be seen in Table 2, these Unbiased Method 5 simulations give hit rates across all six complexes of 32%, 38.9%, 6.2%, and 15.9%, for RMSD_RF, RMSD_RF_PP, ligand_E, and ligand_E_PP, respectively. From these findings, it is tempting to decide that RMSD_RF_PP should be selected as the sole filter of choice for Method 5. However, it can also be seen in Table 2 that the RMSD_RF and RMSD_RF_PP filters yield very small final ensembles for the Unbiased simulations – anywhere from eight ternary complexes for 6HR2_U to a single (yet crystal-like) pose for 6BN7_U and 6HAY_U. While this parsimony might generally be viewed as beneficial – after all, the goal of any method modeling PROTAC-mediated ternary complexes would surely be a single structure that reliably experimental characterization data – until this perfect modeling approach is developed, the danger is that such a restrictive simulation might only provide incorrect results, if it provides any output at all. As will be shown below when various case studies are considered, this expectation unfortunately proves true, and thus all case studied below using Method 5 considered all four filters, despite the better performance for RMSD_RF_PP on the crystallographically characterized validation set.

Finally, given the small ensembles generally produced with Method 5, no overall enrichment could be achieved when the double clustering approaches detailed for Method 4B were applied (data not shown). However, because Method 5 only requires a single (Unbiased) protein-protein docking simulation, and because there is no search of the PROTAC conformational space (as is performed in Methods 2-4B), Method 5 is notably faster than Method 4B, and for this reason alone it warrants further consideration.

## RESULTS

The validation set of ternary complexes utilized above for method development (Table 1) have all been experimentally characterized via X-ray crystallography. However, the overwhelmingly more common scenario is one where PROTAC(s) have been assayed for their ability to degrade a target using a variety of experimental techniques – most commonly with immunoblotting^40^ measured at discrete time points – without any structural elucidation of the ternary complex intermediate. Thus, the outstanding question is whether the computational modeling methods presented earlier^41^ and above can yield insight when no ternary complex crystal structure exists to compare against. In other words, even if the ability of these methods to (retrospectively) reproduce ternary complex crystal structures has been established, does this mean that those methods can predict whether new PROTACs can degrade the target proteins? As will be shown below across seven individual case studies, the answer to this question is generally “Yes.” In particular, it will be shown that the *population of the largest double cluster generated by Method 4B* can be used as a score to distinguish between potent and ineffectual PROTACs. The retrospective case studies below establish this capability across different targets, using a wide variety of PROTAC architectures (*i*.*e*., with different linker compositions, rigidities, lengths, and attachment points to the corresponding binding moieties), and utilizing different E3 ligases – including one, cIAP1,^58^ where no ternary complex crystal structure has yet been solved, and this the methods developed in this work based on VHL- and cereblon-containing complexes may prove less accurate.

### Case Study 1: Evaluation of a New Ternary Complex Crystal Structure of a Macrocyclic PROTAC

However, before we consider systems where no ternary complex crystal structures have been solved, first we consider the accuracy of Method 4B for a very recent ternary complex crystal structure, 6SIS.^56^ (Due to the unique macrocyclic nature of the PROTAC in 6SIS, we did not perform the necessary modifications to adapt Method 5’s linker path finding algorithm to map out two distinct water channels for two different portions of the macrocycle). It should be noted that, although 6SIS was not included in the dataset used to develop Method 4B, this crystal structure strongly resembles one of the crystals in the validation set, 5T35 (*i*.*e*., both contain VHL as the E3 ligase, Brd4^BD2^ as the target, and superpose to one another with a C_α_ RMSD of 0.6Å), and thus it should be expected that Method 4B can effectively model this new crystal structure.

As can be seen in Table 3, Method 4B is indeed very accurate when applied to 6SIS, despite having no (direct) knowledge of the ternary complex crystal structure during the method development phase. Indeed, while the hit rate is 88% using the <10Å RMSD crystal-like criterion, *all* of the modeled poses in the largest double cluster for 6SIS satisfy the CAPRI-like high/medium/acceptable criteria (with 62/184/12 poses, respectively). It is also interesting to note the correspondence between the Method 4B results and various experimental measures of degradation potency, particularly when MZ1, the PROTAC from 5T35, is compared against this new macrocyclic PROTAC. Specifically, Method 4B yields a largest double cluster population for 5T35 roughly 4x larger than for 6SIS, which generally mirrors the experimentally assayed data. Any measured differences between these two systems can be attributed solely to the PROTACs themselves, as both the E3 ligase and degraded targets are otherwise identical. Some of the inherent differences between the two PROTACs are beyond the scope of Method 4B’s ability to model (*e*.*g*., differences in cell permeability), but regardless the results of Method 4B, suggesting that MZ1 should be a more effective degrader than the macrocycle, nevertheless follow the experimental results.

**Table 3.** Comparison of Method 4B Results with Experiment for Two VHL-Brd4^BD2^ Systems.

| PDB | PROTAC | Method 4B<sup>a</sup> | Crystal-like (Hit Rate) | DC50<sup>b</sup> | EC50<sup>c</sup> |
| --- | --- | --- | --- | --- | --- |
| 5T35 | MZ1 | 1040 | 979 (94.1%) | 0-5 | 191 |
| 6SIS | 1 (Macrocycle) | 258 | 227 (88.0%) | 25-125 | 643 |

<sup>a</sup>Population of the largest double cluster <sup>b</sup>In nM, for Brd4(long) <sup>c</sup>In nM, from 22RV1 cells at 72h

### Case Studies 2 and 3: Revisiting Predictions Originally Made with Method 4 – (2) Wild Type vs. Mutant and (3) Three Bromodomains with Three PROTACs

In our original publication on PROTAC modeling,^41^ two systems were put forth to validate Method 4’s performance. For the first (Case Study 2), a triple-point mutation originally proposed by Gadd *et al*.^54^ was modeled. This mutant (dubbed the QVK mutant) was designed based on the 5T35 crystal structure to force the original target Brd4^BD2^ protein to more closely resemble Brd2^BD1^, a protein less effectively degraded by the PROTAC MZ1. The results generated by Method 4^41^ correctly gave fewer final ternary complexes for the mutant than for the wild type, regardless of the particular set of filters and thresholds used (*i*.*e*., 4^A^ to 4^F^), in agreement with experiment.

For Method 4B – which is not to be confused with Method 4 using the “B” set of filtering criteria in Ref. 41, *i*.*e*. 4^B^), as mentioned above the most effective scoring function is the population of the largest double cluster. With Method 4B, this population was found to be 1040 for the wild type (Table S1) and 124 for the QVK mutant – again in agreement with the experimental result. The corresponding predictions from Method 5 can be found in Table S2 using all four variants; only ligand_E_PP fails to correctly follow the diminished degradation for the QVK mutant.

The other validation system considered in our original publication (Case Study 3) involves an analysis of the work of Zengerle *et al*.,^60^ who investigated the degradation of three similar but distinct targets – Brd2, Brd3, and Brd4 – using three similar but distinct PROTACs – MZ1, MZ2, and MZ3. In general, they found that the PROTACs decreased in degradation potency from MZ1>MZ2>MZ3, and similarly that the targets were selectively degraded in the order Brd4>Brd3>Brd2. Using the same target crystal structures as in our previous work^41^ for modeling inputs (5T35, 3ONI, and 3S92 for the second BDs of Brd4, Brd3, and Brd2, respectively), both Method 4B and Method 5 were used to model whether these experimental orderings could be (retrospectively) predicted; the results are shown in Tables 4 and S3, respectively.

**Table 4.**
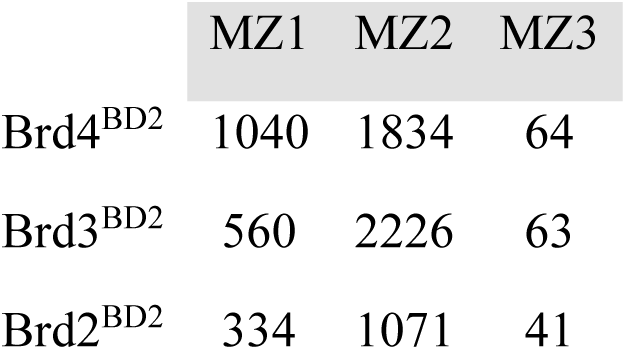
Predictions with Method 4B for Three PROTACs on Three Brd Targets (Case Study 3)

If the results in Table 4 perfectly matched the experimentally assays of Zengerle *et al*.,^60^ the double cluster populations would decrease from both left-to-right across a row (thereby indicating MZ1>MZ2>MZ3) and top-to-bottom down a column (for Brd4>Brd3>Brd2). While such agreement can indeed be noted in some cases – down the MZ1 and MZ3 columns, for instance – it seems that the degradation efficacy of MZ2 is consistently overpredicted by Method 4B. MZ2 differs from MZ1 in that it has four, rather than three, PEG units in its linker, and thus is longer and more flexible than MZ1. Correspondingly, the rigid binder conformational search of Method 4B generated 203 initial PROTAC conformations for MZ1, but 1308 for MZ2, a 5x increase. While the filters and clustering procedures incorporated into Method 4B ultimately rendered many of these additional input conformations ill-suited to form ternary complexes – note (Table 4) the 1.8x, 4.0x, and 3.2x increase from MZ1 to MZ2 for Brd4, Brd3, and Brd2, respectively, rather than the 5x difference in the number of input PROTAC conformations – Method 4B nonetheless generally does seem to favor the greater flexibility afforded by the larger linker in MZ2. A recent preprint^43^ has accounted for a similar phenomenon by normalizing their final results based on the initial number of PROTAC conformations (albeit with an artificial maximum of 1000 conformations allowed). We have instead opted to present our results as-is, without any such normalization, as these unadjusted results still generally match the experimental findings quite well, as will be shown in the rest of the case studies in this work. Nonetheless, for this particular case study, it can be fairly observed that simply increasing the flexibility artificially led to an increase in the predicted efficacy of a PROTAC. This phenomenon is thus analogous to how merely increasing the size of a small molecule inhibitor can often artificially increase its predicted binding affinity.^61^ In any event, the impact of the conformational flexibility of a PROTAC on its ability to degrade its target will be revisited in the additional case studies considered below. In the meantime, it is interesting to note that Method 5 – which does *not* utilize a PROTAC conformational search as part of its methodology – is somewhat more effective at predicting the correct PROTAC ordering (as evidenced by the frequent decreases from left-to-right in Table S3), even if the proper ordering for the degradability of the target proteins is never correctly predicted (*i*.*e*., the data never decreases from top-to-bottom in Table S3).

### Case Study 4: The Ability to Model Cereblon-recruiting PROTACs

As can be seen in Table 1, Table S1, and Figure 3, Method 4B does a much better job reproducing known ternary complex crystal structures when the E3 ligase is VHL rather than cereblon; Method 5 does not exhibit this disparity quite so starkly. It is therefore necessary to fully explore how well cereblon-mediated TPD can be modeled, particularly using Method 4B. To that end, for Case Study 4 we consider the system experimentally characterized by Zorba *et al*.,^48^ where the kinase BTK was successfully degraded using an ibrutinib-like binding moiety linked to the cereblon binding moiety pomalidomide via a series of 11 PEG-based linkers of varying lengths. It should be noted that studies such as this one – with one target and one E3 ligase but a library of possible PROTAC linkers – are the most common situations explored in the peer-reviewed TPD literature.

Experimentally,^48^ Western blots of cell lysates showed that the two shortest PROTACs (1 & 2) were unable to degrade BTK, the next two shortest (3 & 4) were marginally effective at PROTAC concentrations >1µM, the next longest PROTAC (5) was moderately effective, and the longest five PROTACs (6-11) were all essentially equally potent at the measured concentrations. A TR-FRET-based assay was also built to directly probe ternary complex formation, which provided results generally mirroring the degradation results; modeled results also correctly found a correlation between linker length and degradation. Importantly, it was also established that, similar to other cereblon-based degraders,^55^ effective target degradation did not require strong or cooperative direct protein-protein interactions. Given that all of the VHL-based ternary complexes used to develop our computational methods do indeed exhibit positive cooperativity, this BTK-cereblon degradation case study should therefore provide a strict test as to whether favorable protein-protein interactions are required for successful predictions.

Figure 4 presents the modeling results for Case Study 4 using Methods 4B and 5 (for the latter, only the ligand_E variant yielded a curve qualitatively similar to the experimental results). As can be seen, Method 4B matches the experimental trends quite well, particularly in capturing how PROTAC 5 is the point after which degradation truly starts to be observed (*i*.*e*., anything shorter than PROTAC 5 is largely ineffective, while anything longer is efficacious). The ligand_E variant of Method 5 instead identifies PROTAC 3 as this “tipping point,” although it should be noted that the observed saturation of degradation efficiency for the longest linkers is better described using Method 5 (even though it is somewhat evident in the Method 4B results).

**Figure 4.**
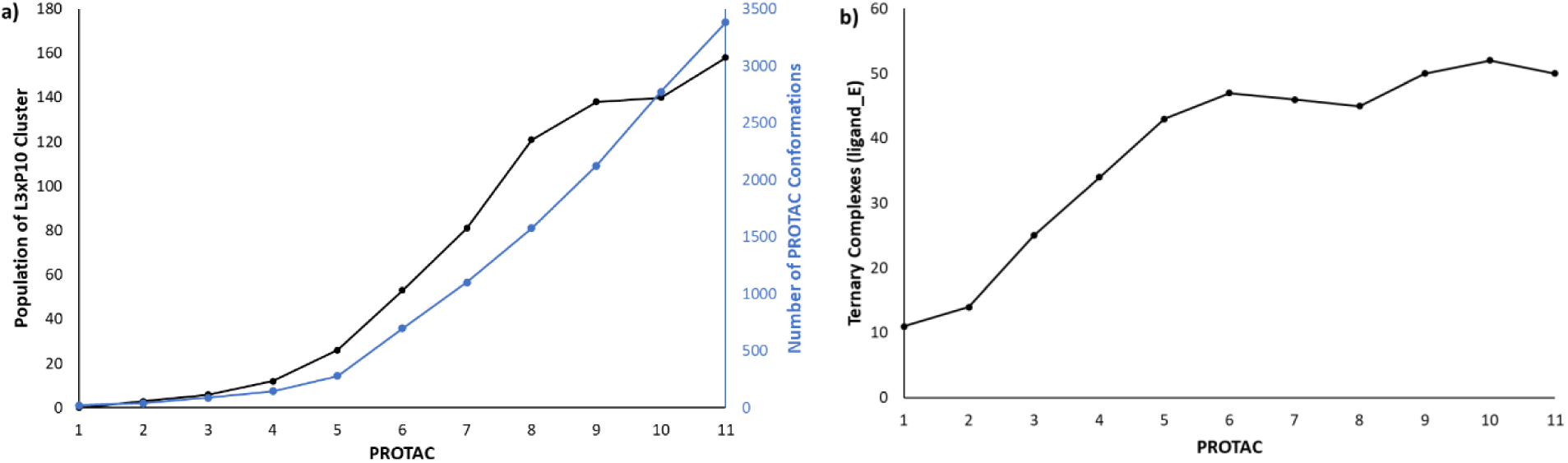
Modeling results for Case Study 4, the BTK-cereblon-PROTAC system of Zorba *et al*.^48^ a) The results from Method 4B: (black) the population of the largest double cluster as a function of the PROTAC and (blue) the number of conformations produced by Method 4B for each PROTAC. B) The results from Method 5, using the ligand_E filtering criteria.

Also shown in Figure 4a is a plot of the number of PROTAC conformations (blue curve) produced with Method 4B. As can be seen, this curve tends to follow the double cluster population plot in black, although not exactly, as the modeled degradation ability seems to flatten out after PROTAC 8, even while the number of PROTAC conformations continue to increase linearly (with a slope of 517 from PROTAC 5 to 11 and an r^2^ of 0.99). Nonetheless, the results shown in Figure 4 do suggest that the modeling tools of this work can successfully model cereblon-containing PROTACs.

Both Methods 4B and 5 require the two (binary) protein-ligand structures as input for protein-protein docking. In Case Study 4, for the E3 ligase+binder, the crystal structure 4CI3 – cereblon liganded by pomalidomide, the exact binding moiety utilized in PROTACs 1-11 – was used as-is, subject only to the routine preparation required to generate partial charges and model in missing hydrogens, sidechains, and short loops unresolved in the crystal structure. However, for the target+binder component, the BTK binding moiety utilized by PROTACs 1-11, while similar to the covalent binder ibrutinib, does differ: specifically, this moiety contains two fluorines on the terminal phenyl ring not found in ibrutinib, and ibrutinib’s aminopyrazolopyrimidine core has been replaced with a monocylic pyrazole core as well. Thus, to generate the BTK+binder binary structure needed for protein-protein docking, the actual target binding moiety of Zorba *et al*.^48^ was docked into the ibrutinib binding site of the crystal structure 5P9I using MOE. The resulting best scoring pose closely resembled ibrutinib’s crystallographic positioning, and thus this docked pose was used to position the binding moiety for the target+binder binary complex.

However, for many protein targets of interest to PROTAC-mediated TPD, there may not exist a cocrystal structure of the target with the desired target binding moiety – or even with a ligand closely resembling the target binding moiety, as was used for Case Study 4, particularly if weak binders are to be incorporated into a putative PROTAC. Below, we will investigate systems where the necessary starting experimental structural information is incomplete, meaning that the input binary structures *themselves* must be modeled, *e*.*g*. via docking or through direct *in situ* modification of a closely related ligand in a pocket. The impact of how this initial modeling is performed will be thoroughly investigated, as detailed below.

### Case Study 5: One PROTAC against Multiple Targets, with no Exact Target+Binder Crystal Structures

Huang *et al*.^62^ utilized a quantitative proteomics approach to identify proteins degraded after exposure to the PROTAC TL12-186, which deliberately incorporates the pan-kinase inhibitor TL13-87 as its target binding moiety (Figure 5a&b). As a consequence of the promiscuity of the chosen binder, TL12-186 was found to bind to many proteins, in particular (and as expected) kinases. However, importantly it was also determined that – even upon considering just the kinases interacting with TL12-186 – PROTAC binding in and of itself was insufficient to ultimately effect TPD, suggesting that “additional factors” contribute to whether a kinase bound to TL12-186 is degraded.

**Figure 5.**
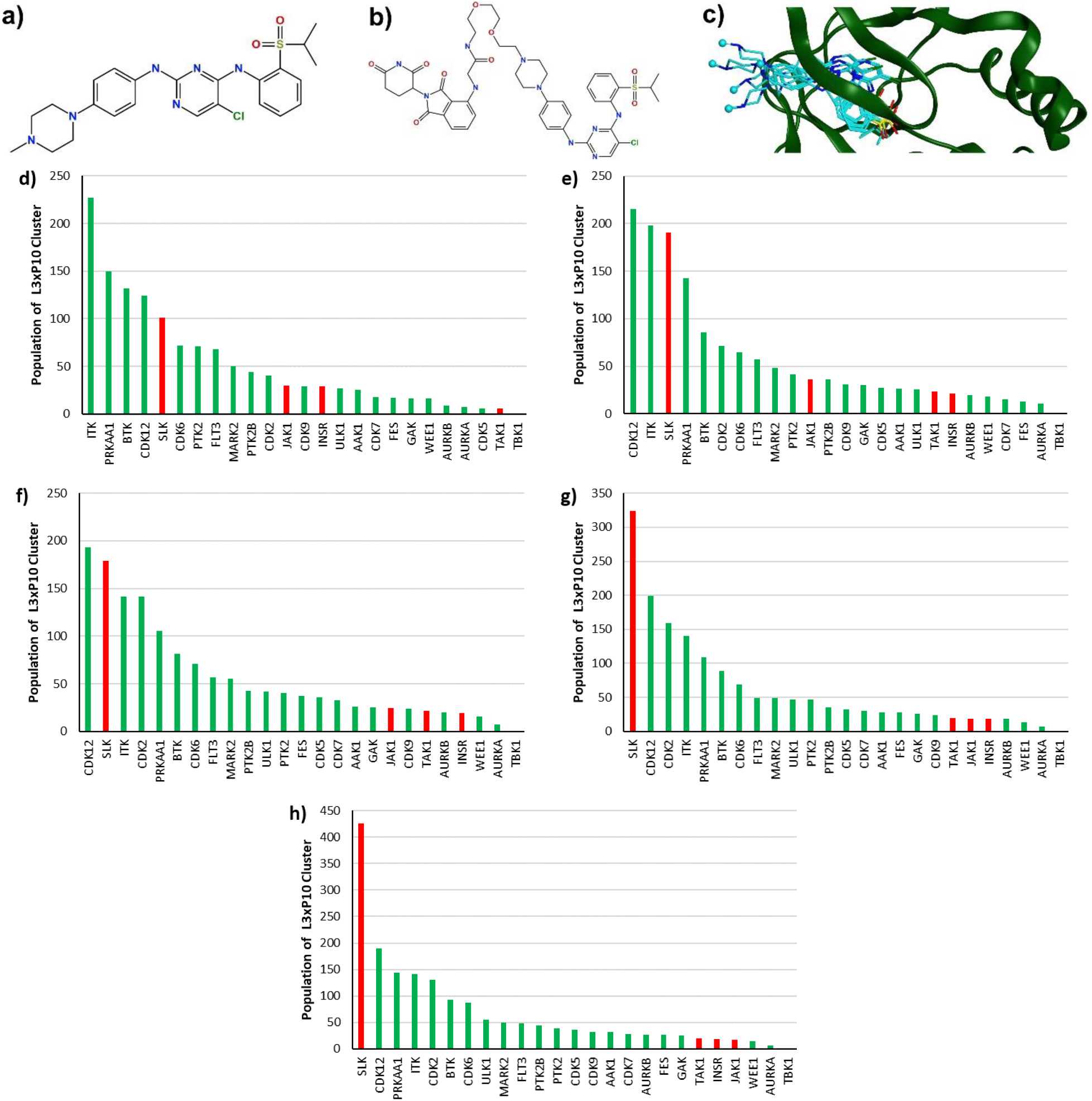
a) Structure of TL13-87, the pan-kinase binding moiety used in b) TL12-186, the multi-kinase targeting PROTAC designed by Huang *et al*.^62^ c) Illustration of the variety in exit vectors across five poses of TL13-87 docked into AURKB; the methyl connection points to the linker of TL12-186 are rendered as cyan spheres. d)-h) The average population of the largest double clusters for Method 4B for each kinase (*x*-axis) averaged across the top d) 1, e) 2, f) 3, g) 4, and h) 5 binary kinase+binder complexes as input. Green bars indicate those kinases found by Huang *et al*. to be degraded by TL12-186, whereas red bars indicate the five kinases found to bind to this PROTAC without subsequent degradation. TBK1, with a population of zero in d)-h), is a non-degraded (red) kinase.

We sought to apply Methods 4B and 5 to this system, fully aware that a tight focus on one particular entity, namely the intermediate ternary complex, ignores many other factors that can influence the results of experimental cellular assays. For Case Study 5, the kinases identified in Table 1 of Huang *et al*.^62^ as being significantly degraded by TL12-186 were considered, as were the kinases indicated in the main body of that text as being “substantially engaged by TL12-186” but that lacked accompanying degradation. To assemble the inputs for this modeling study, we first applied MOE’s Project Search capability to the MOE Protein Kinase database of annotated kinase crystal structures, to identify potential target+binder binary complexes that could be used as initial inputs for the protein-protein docking steps of Methods 4B and 5. However, no public crystal structures were found for any of the selected kinases that were also cocrystallized with the binding moiety TL13-87. Thus, alternatives were manually selected – based partly on the overall (Tanimoto) similarity between the actual cocrystallized ligand and TL13-87, but mostly based on whether the cocrystallized ligand possessed a solvent-exposed group analogous to the methylpiperazine group of TL13-87 (which would indicate a possible exit vector/attachment point for the rest of the PROTAC). The initial binary kinase crystal structures ultimately chosen are listed in Table S4; Figure S2a also provides the Tanimoto similarities between TL13-87 and these cocrystallized ligands, which ranges from 0.87 (FES) to 0.31 (GAK).

After these initial kinase crystal structures were identified, it was then necessary to dock the actual binding moiety (TL13-87) into each kinase pocket. Although docking is a routine structure-based drug design technique, it was *a priori* uncertain if a single docked pose for TL13-87 with each kinase could accurately represent the positioning of the kinase binding moiety in a putative ternary complex, particularly as the rest of the PROTAC is known to interact with the binding moiety.^54^ Even a subtly different placement of the solvent-exposed methylpiperazine connection point of TL13-87 could potentially alter not only the protein-protein interaction landscape explored by protein-protein docking, but also the orientation of the linker of PROTAC TL12-186. For example, if an initial kinase+binder binary pose oriented its terminal methyl connection point towards a wall of the kinase, this geometry would sterically preclude the extension of TL13-87 out to the rest of the PROTAC. We therefore sought to identify the best way to account for the variability imposed by the initial target+binder positioning, as measured by the accuracy of the final predicted results, *i*.*e*., the relative degradation of the kinases in Table S4.

To build various binary target+binder inputs, each of the 25 kinase crystal structures tabulated in Table S4 – 20 of which degrade after PROTAC binding – was superposed onto PDB 2XB7 (the crystal structure originally used by Huang *et al*.^62^ to design their TL13-87 pan-kinase binding moiety). The position of the native ligand in 2XB7 was then directly transferred over to the respective pockets in each of the 25 studied kinases, thereby roughly outlining a potential initial binding orientation for TL13-87. From this starting point, MOE Dock was used to more rigorously and systematically optimize the rough placement of TL13-87 in the binding pockets using two different approaches: General docking (*i*.*e*., the default non-covalent docking approach) and Template-based docking, where the crystallographic orientation of the chloropyrimidine-amino-phenylsulfonyl core copied over from 2XB7 was used to initially place the matching central scaffold of TL13-87, after which point potential docking poses were relaxed in a rigid receptor pocket prior to scoring. Up to 30 (non-duplicate) poses were generated with each docking approach and were scored using the default GBVI/WSA dG scoring function.^63^ These two ensembles of docked poses were then combined, sorted by docking score, and finally inspected, with five unique poses manually selected as the final input structures for each of the 25 binary kinase+binder complexes. These five poses were chosen so that a) the methylpiperazine of TL13-87 extended out of the kinase binding pocket and into solvent, while allowing sufficient space (judged visually) to connect to the linker and cereblon binder of TL12-186, b) no duplicate poses were included, and c) poses with better (more negative) GBVI/WSA dG docking scores were preferred. As an example of the final result, Figure 5c depicts the five input poses for AURKB+TL13-87 generated by this procedure. As can be seen, while the core deep in the pocket varies only slightly, the presented exit vector of the methylpiperazine is placed in five different positions.

Armed with a collection of varied target+binder poses for each kinase, protein-protein docking was then used to separately dock each of these five kinase+binding poses for each kinase against the prepared cocrystal structure of cereblon and pomalidomide (as was used in Case Study 4). For Method 4B, a conformational ensemble of the PROTAC TL12-186 was generated once and was then used for the result of the simulations. Figures 5d-h show the results for Method 4B, where the populations of the largest double cluster were averaged for each kinase across the top, top 2, top 3, etc. ranking input poses (as scored by GBVI/WSA dG). In these plots, the predicted degradation decreases from left to right, and the red bars indicate the five kinases (INSR, JAK1, SLK, TAK1, and TBK1) found to *not* degrade by Huang *et al*.^62^ (Taking the median of the top *N* double cluster populations instead of the average gives similar results; data not shown). Thus, a simulation that perfectly identifies the non-degrading kinases (ignoring considerations of the cellular environment not accounted for by Method 4B, as previously mentioned) would show five smaller red bars on the far-right of the plots of Figures 5d-h. As can be seen in Figure 5d, considering just the single top-scoring kinase+binder input pose for each kinase shows little discrimination between the degrading kinases (green) and the non-degrading kinases (red). However, as more input kinase+binder poses are considered, three of the non-degrading kinases (TAK1, INSR, and JAK1) slide to the right of the plots to join with TBK1 on the far-right, indicating that these four kinases that were experimentally found to *not* be degraded by TL12-186 are correctly predicted to be less and less degraded, if additional input poses are considered. By contrast, however, the fifth non-degraded kinase, SLK, is predicted to be *more* degraded as additional input poses are considered, to the extent that it is ultimately predicted to be the kinase *most* susceptible to degradation by TL12-186 if the top 5 input poses are averaged.

One possible explanation for this misprediction is that the GBVI/WSA dG binary docking scores (Figure S2c) for TL13-87 with SLK are the lowest on average, and so the modeled degradation shown in Figure 5 may be offset by generally poor binding between SLK and TL12-186. However, Huang *et al*.^62^ noted no such corresponding phenomenon in their experiments; moreover, overall there is no apparent correlation overall between the average binary docking scores and the red or green kinases of Figure 5. Another possible explanation stems from early observations made when Method 4 rather than 4B was applied to this system. For Method 4, ternary complexes are included in the final output only if they have at least 100Å^2^ of buried hydrophobic surface area at the protein-protein interface. With this requirement (Figure S2d), SLK scores considerably more poorly (although it is still the best among the five non-degraded kinases). Thus, it may be that the move to the single “one-size-fits-all” filtering criteria in Method 4B led to poorer results in this specific instance. The findings of Method 5 also support this hypothesis, as the ligand_E_PP variant of Method 5 – which considers the protein-protein interface – also overpredicts the degradation of SLK (Figure S2e). By contrast, the RMSD_RF variant of Method 5 (Figure S2f), where there is no such consideration of the protein-protein interface, gives the best overall predictions for the four variants of Method 5, correctly ranking SLK near the bottom of all kinases.

All in all, with the single exception of SLK for Method 4B, the performance exhibited by both Methods 4B (Figure 5) and 5 (Figure S2) for Case Study 5 is certainly encouraging. However, in order to obtain the most accurate predictions, it seems *multiple possible input poses must be considered* if the target+binder complex geometry is particularly flexible or is otherwise ambiguous, *i*.*e*. it has not explicitly been characterized with X-ray crystallography. It seems likely that other approaches, such as molecular dynamics sampling of the input binary structures (or of each generated ternary complex), would also similarly be capable of delivering this improved accuracy. In a sense, the static docking-based workflow presented above represents discrete snapshots (five, in this case) of an evolving dynamic trajectory. In any event, the improved accuracy resulting by considering multiple possible input orientations will also be utilized in the modeling results of the next two case studies.

### Case Study 6: Multiple PROTACs against Multiple Targets, with no Exact Target+Binder Crystal Structures

For Case Study 6, we return to the common scenario where multiple PROTACs are evaluated for their abilities to degrade their targets – in this case, these targets are the similar but distinct BDs found in three BET (bromo- and extra-terminal) proteins degraded by the PROTAC library designed by Qin *et al*.^64^ Based on a novel [1,4]oxazepine target binder (QCA276), many of the PROTACs explored in this work were shown to be effective degraders, even at low picomolar concentrations, as evaluated in multiple human leukemia cell lines and for tumor xenografted mice. Moreover, the PROTAC linkers utilized by Qin *et al*.^64^ are generally more rigid than the PEG- or alkyl-based linkers typically utilized during initial PROTAC development. Thus, the PROTACs of Case Study 6 also often incorporate an alkynyl moiety in their linkers, as well as an alkynyl-pyrazole moiety that, while nominally part of the BET binder, could alternatively be viewed as a rigid spacer installed at one end of the linker (Figure 6a).

**Figure 6.**
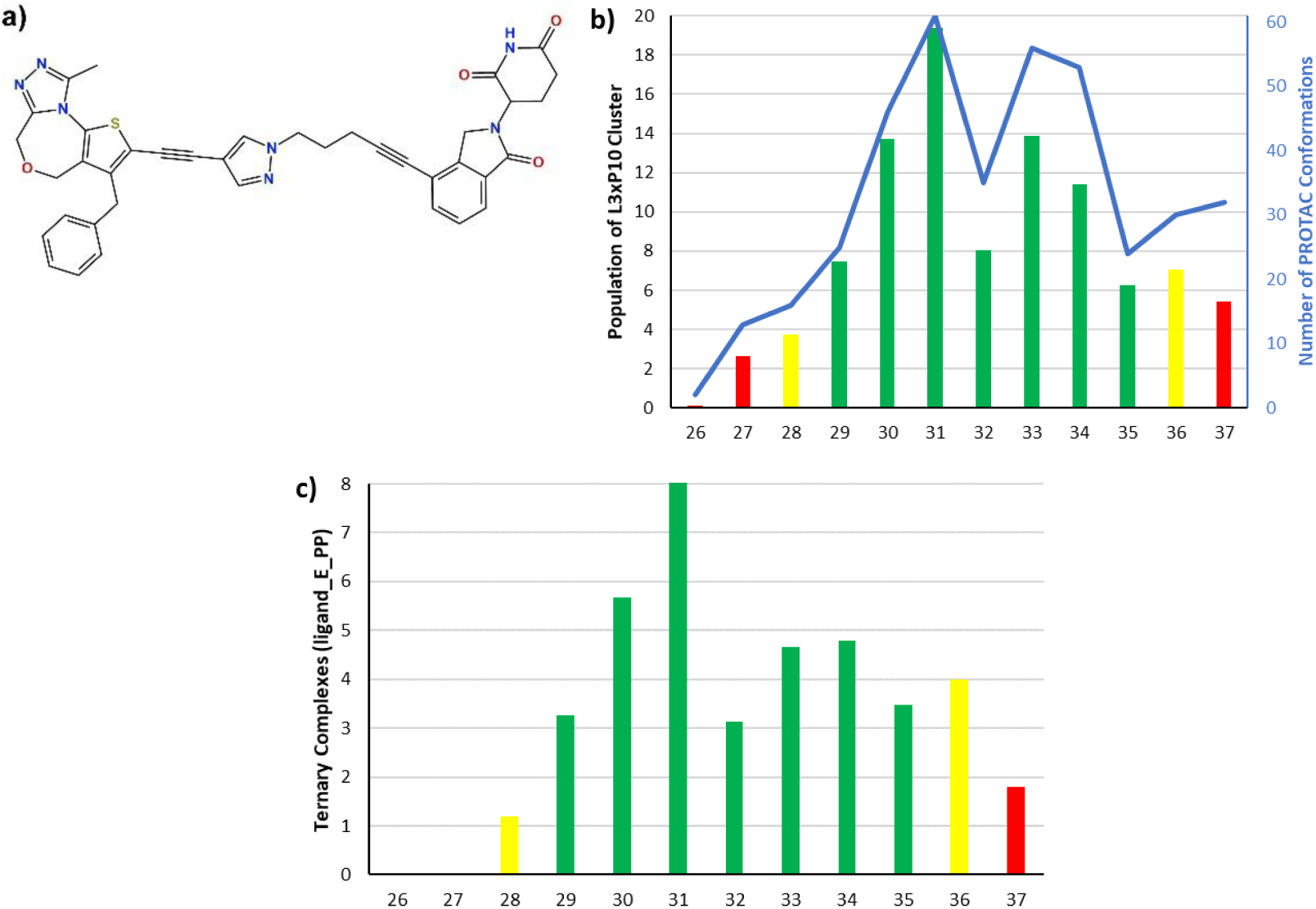
a) Structure of Compound 35 (QCA570), a representative PROTAC from Qin *et al*.^64^ b) Method 4B modeling results for Case Study 6, averaging the double cluster populations across multiple input poses for Brd2^BD2^, Brd3^BD2^, and Brd4^BD2^. The green, yellow, and red bars correspond to PROTACs with high, intermediate, and low experimentally assessed IC50s. Also shown (blue line) is the number of conformations for each PROTAC generated with Method 4B. c) Method 5 modeling results for Case Study 6, using the ligand_E_PP variant, with the same green/yellow/red scheme used in b).

The QCA276 target binder designed by Qin *et al*.,^64^ although developed based on a structure of Brd4^BD1^ in complex with the pan-BET inhibitor JQ1 (which was also incorporated as the binding moiety of PROTAC MZ1 cocrystallized in the ternary complex crystal structure 5T35^54^), was measured to bind at nanomolar concentrations to both the first *and* second BDs of Brd2, Brd3, and Brd4. Crystal structures exist for all six BDs, although none with the exact QCA276 binding moiety. However, considering the parent JQ1 binder, cocrystal structures exist for five of the six possible combinations, *i*.*e*. for all except Brd2^BD1^. Thus, in assembling the inputs for the binary target+QCA276 components for this case study, it was decided to model only the three binary complexes of QCA276 with the second BDs of Brd2, Brd3, and Brd4, as it was deemed that the missing starting point for Brd2^BD1^+JQ1 might unduly impact the simulations.

Therefore, the same binary docking protocol detailed above (Case Study 5) was applied to generate the input for modeling Qin *et al*’s work^64^. Specifically, QCA276 was docked into the JQ1 binding site of the BD2 proteins listed in Table 4 (*i*.*e*., 3ONI for Brd2^BD2^, 3S92 for Brd3^BD2^, and 5T35 for Brd4^BD2^). As before, both General and Template-based docking protocols were applied to generate diverse starting orientations for the binder. In addition, for each protein, the original, crystallographically resolved JQ1 was transformed into QCA276 directly in the pocket via manual ligand modification followed by a minimization of QCA276 with neighboring protein residues restrained via tethers to stay near their original positions. These *in situ* derived BET+QCA276 complexes were always considered as inputs, as were the top docked poses selected using the same criteria described earlier in Case Study 5. For Brd2^BD2^, only three unique docking poses of QCA276 were generated, and thus four total poses (including the binary complex generated via *in situ* ligand modification) were used as inputs for the protein-protein docking phase.

For the E3 ligase components, the PROTAC library considered in this case study (consisting of compounds 26-37 of Qin *et al*.^64^) utilized cereblon with a variety of different binding moieties: thalidomide for compounds 26, 27, and 36; pomalidomide for 28-31; and lenalidomide for 32 and 33. Additionally, some compounds utilized derivatives of these three parent cereblon binders: *e*.*g*. 34 and 35 used lenalidomide without the amino group connected to the linker, and 37 incorporated a methylated lenalidomide. However, inspection of the different cereblon+binder crystal structures (*e*.*g*., 4TZ4 for cereblon with lenalidomide, 4CI3 with pomalidomide [as used in Case Studies 4 and 5], and 4CI1 with thalidomide) revealed negligible structural differences in the binding sites, and thus 4TZ4 was (arbitrarily) chosen as the sole representation of cereblon, with the different binding moieties utilized in the PROTAC library modeled *in situ* (as was done for the target+QCA276 binary complexes) and described with only a single, crystallographically-informed pose.

The 11 PROTACs of this case study can be qualitatively grouped into three classes based on their IC_50_ values measured across the three leukemia cell lines considered by Qin *et al*.^64^: for example, considering the RS4;11 line, the most potent PROTACs (compounds 29-35) all show IC_50_ values between 0.0012 and 0.038 nM; the intermediate group (compounds 28 and 36) are an order of magnitude less potent, although still subnanomolar (0.379-0.39 nM); and the final group (compounds 26, 27, and 37) are two or more orders of magnitude less potent (20.8-1000+nM) still. These three categories have been colored as green, yellow, and red, respectively, in Figure 6, where the modeling results using Method 4B and Method 5 (with the ligand_E_PP variant) are presented. As can be seen, Method 4B separates the 11 PROTACs into their corresponding experimental categories quite well: among the degraders experimentally determined to be the best (green), only compound 35 is predicted by Method 4B to be less effective than any of the intermediate (yellow) or low (red) tier PROTACs. The performance using Method 5 is also satisfactory, if not quite as solid as that shown for Method 4B: specifically, compound 36 (yellow) is predicted to outperform three of the most effective compounds (39, 32, and 35). Regardless, both methods would suggest deprioritization of compounds 26, 27, 28, and likely 37, which nicely mirrors the experimental findings.

Also shown in Figure 6b (blue) are the number of initial conformations generated for each PROTAC using Method 4B. As was shown in Case Study 4 (Figure 4a), there is a correspondence between this raw number of PROTAC conformations and the final number of ternary complexes found in the most populous double cluster (as averaged across multiple target+binder input complexes). However, as was also seen in Case Study 4, this correlation is modulated by the ability of the conformations in each ensemble to effectively connect the two pockets in the protein-protein docked ensemble. For example, based strictly on the number of conformations, compounds 36 (yellow) and 37 (red) would be predicted to be more effective degraders than compounds 29 and 35 (green), whereas the actual final Method 4B results rank 29 highest among these four, and also place 35 over 37. Regardless, it is interesting to note that the raw number of generated PROTAC conformations correlates fairly well with the overall efficacy of these PROTACs, even for these highly optimized and highly potent compounds.

### Case Study 7: Degradation Catalyzed by cIAP1, a non-VHL/non-cereblon E3 Ligase

Although nearly 10 E3 ligases have been exploited in a TPD context,^3,19^ only two – VHL and cereblon – have been structurally characterized via X-ray crystallography in an intermediate ternary complex. Furthermore, although to date VHL and cereblon are undoubtedly the two E3 ligases most commonly utilized for TPD,^23^ alternatives are highly sought, both because E3 ligase expression levels may vary in different tissue or cellular environments^21^ and as a potential means to combat evolved resistance to PROTAC-mediated TPD.^17^ Beyond VHL and cereblon, the third-most commonly utilized E3 ligases in a TPD context are the inhibitor of apoptosis (IAP) proteins. Indeed, the subclass of PROTACs incorporating an IAP-recruiting moiety are often termed SNIPERs (specific and nongenetic inhibitor of apoptosis protein-dependent protein erasers) by their developers. We were interested to determine whether Methods 4B and 5, which were developed based on PROTAC-mediated ternary complex crystal structures utilizing only VHL and cereblon, would be capable of modeling TPD mediated by a different E3 ligase.

For this case study, we selected the work of Shibata *et al*.,^58^ who developed extensive libraries of SNIPERs, many of which were shown to be effective in degrading their target protein, the androgen receptor (AR). As part of their work, every component of their SNIPERs was explored, including two different binding moieties for cIAP1 (the specific IAP E3 ligase utilized), multiple variants of the AR binding moiety, and linkers of varying lengths, compositions, and attachment points to the two respective binding moieties (see Table S5 for a list of all SNIPERs considered in Case Study 7). The structure of PDB 1Z95, the AR ligand binding domain in complex with bicalutamide, was chosen to represent the target protein, as this structure was the one considered by Shibata *et al*.^58^ to determine ideal linker attachment points. The unique target binding moieties found in the library of SNIPERs (Table S5) were docked into the bicalutamide binding pocket of 1Z95 using MOE’s Template-based docking approach, with potential poses generated by matching the common cyanophenyl cores, followed by pose minimization in a rigid receptor pocket. Due to the tight constraints of the pocket, only a single representative pose was selected for each unique binding moiety (*cf*. the target pose generation procedure of Case Studies 5 and 6); the selected poses across those different moieties nonetheless all superpose closely, as shown in Figure S3a.

To represent the input cIAP1+binder structures, three different cIAP1 crystal structures (3MUP, 4HY4, and 4KMN) were considered as initial inputs, consisting of the BIR3 domain of cIAP1 cocrystallized with different previously designed small molecule mimetics of the endogenous tetrapeptide ligand of cIAP1. As can be seen from Figure S3b, these three possible structures, although structurally similar in general, differ with regard to the placement of their C-termini. For this reason, the two cIAP1 binders incorporated into the SNIPERs of Shibata *et al*.^58^– bestatin and compound 24 – were docked into the pockets of these three cIAP1 crystal structures using MOE’s General Docking procedure. As was done in previous Case Studies, five poses were manually selected for each of the two cIAP1 binding moieties, and across each of the three cIAP1 orientations, yielding a total of 15 input poses for bestatin+cIAP1 and 15 for compound 24+cIAP1, differing in both the placement of the cIAP1 C-terminus and by the orientation of the atom where the SNIPER linkers are attached (see Figure S3c for a representative example).

Methods 4B and 5 were applied to combine these various inputs with the designated SNIPERs of Shibata *et al*.^58^ to form predicted ternary complexes. Whenever possible, results common to multiple simulations were reutilized rather than regenerated *de novo*. For example, all SNIPERs of number 31 and higher vary only in their linker, and contain both the same AR binder (as used in SNIPER-2) and the same cIAP1 binder (compound 24). Thus, the separate protein-protein docked ensembles generated by docking 1Z95 with the binder of SNIPER-2 against 3MUP, 4HY4, and 4KMN and their five bound poses each for the compound 24 binder were reused for all SNIPERs that contained the cIAP1 binding moiety 24. The protein-protein docked ensembles built with Method 4B were also reused for Method 5. However, overall Method 5 failed to give usable results, often failing to produce any output ternary complexes, regardless of the particular variant used. This failure can likely be attributed to the deeply buried AR binding pocket of 1Z95, which often frustrated the ability of the solvation-based path finding algorithm to locate an unobstructed path for the linkers to follow while spanning between the two binding pockets.

For Method 4B, the results presented below were averaged across all possible 15 cIAP1 + binder orientations docked against the single corresponding AR + target binder pose; averaging individually across only a single set of cIAP1 inputs (*e*.*g*., those produced only using 3MUP) does change the precise predicted ordering of SNIPER efficacy, but does *not* change the qualitative results described below and shown in Figure 7. The difference between SNIPERs utilizing the non-specific bestatin as a cIAP1 recruiting moiety vs. the more optimized binder of compound 24 is shown in Figure 7a, where the SNIPERs have been ordered along the *x*-axis (increasing left-to-right) by their experimentally measured efficacies, with a corresponding green (effective) *vs*. red (ineffective) coloring scheme as well. In the experimental assays (Table S5), treatment with 30 µM of the bestatin-incorporating SNIPERs leaves anywhere from 26% (SNIPER-23) to 88% (SNIPER-27) of AR undegraded. Clearly, the efficacies as predicted by the (average) populations of the largest double clusters do not match the experimental trends very well. More satisfying, however, is the finding that SNIPER-31, which differs from SNIPER-2 only insofar as bestatin has been replaced by compound 24, is correctly predicted to be much a more effective degrader than SNIPER-2. Experimentally, at 3 µM, these two SNIPERs leave 30% and 96% of AR undegraded, respectively, while Method 4B predicts an average largest double cluster population of 121 for SNIPER-31 and only 9 for the less effective SNIPER-2.

**Figure 7.**
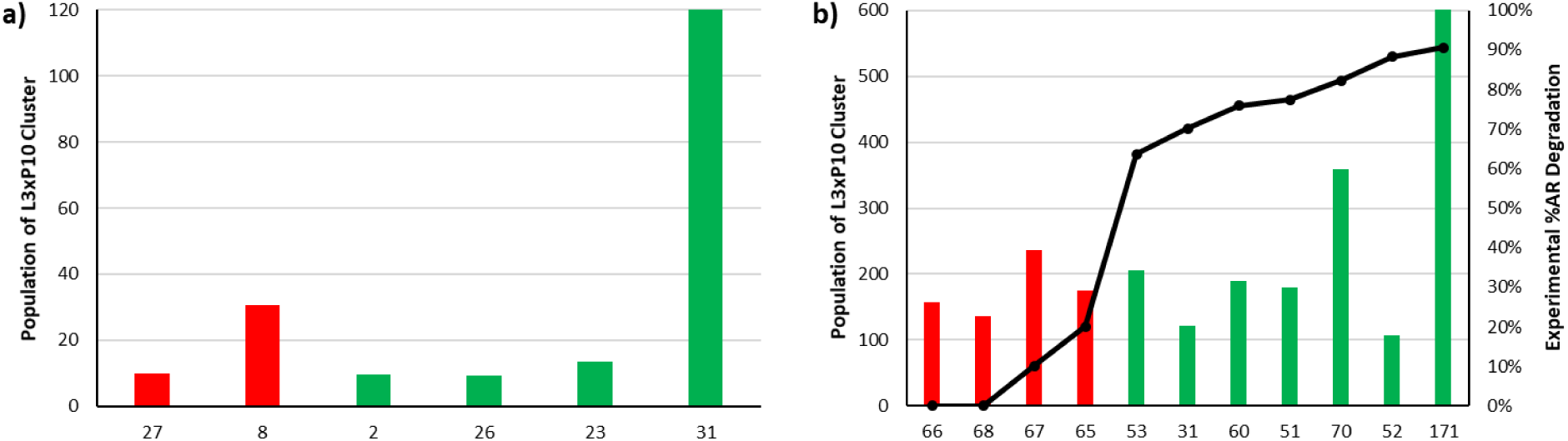
a) Method 4B modeling results for SNIPERs with bestatin (first five bars) or compound 24 (SNIPER-31) as the cIAP1 recruiting moiety. The *x*-axis gives the SNIPER number, sorted from left-to-right by increasing experimental efficacy (see Table S5), and colored red or green for less or more effective SNIPERs, respectively. b) Method 4B modeling results and experimental %AR degradation at 3µM (black line, right axis) for SNIPERs using compound 24 as their cIAP1 binder. Bars are colored and sorted as in a).

However, a comparison of the SNIPERs that exclusively utilize compound 24 as the cIAP1-recruiting moiety (Figure 7b) again reveals underwhelming performance for Method 4B, particularly when compared to the accurate results demonstrated in the earlier case studies of this work. The black line in Figure 7b shows the %AR degraded, with the red bars for SNIPERs 65-68 indicating that these compounds should have lower populations for their largest double clusters. Considering the overall rank-ordering of the SNIPERs in Figure 7b, three of these less-effective degraders (65, 66, and 68) are indeed predicted to be among the five worst SNIPERs, but the fourth experimentally ineffective degrader (SNIPER-67) is predicted to be effective than it was measured to be. Furthermore, although SNIPER-171 is correctly predicted to be the most effective degrader constructed by Shibata *et al*.,^58^ the *second*-most effective degrader, SNIPER-52, is predicted by Method 4B to be the *least* effective SNIPER of those predicted in Figure 7b.

In summary, it appears that although some qualitative trends were correctly recapitulated with Method 4B – particularly the superior performance of SNIPERs incorporating compound 24 vis-à-vis bestatin, the high efficacy of SNIPER-171, and (somewhat) the relative disfavoring of three of the four less effective SNIPERs – the overall quality of these predictions is clearly diminished as compared to the results of the earlier case studies, where degradation was effected by the same two E3 ligases – VHL and cereblon – that were used to construct Method 4B.

## DISCUSSION AND CONCLUSION

The various modeling methods we have developed – including Methods 1-4 of our previous work^41^ and the new Methods 4B and 5 now presented – were developed based on properties observed for a handful of crystal structures of PROTAC-mediated ternary complexes. However, despite this limited training set (which includes only two members out of over 600 possible E3 ligases, as well as structurally similar bromodomain targets), these computational methods – particularly Method 4B – prove well-capable of reproducing experimental findings across a variety of scenarios. As a result of this accuracy, ongoing PROTAC discovery efforts can be aided by a computational modeling component, thereby reducing both cost and time to discovery. The modeling results presented herein are robust as well, even for scenarios where it might be expected that modeling could encounter some difficulties, such as when a uniquely macrocyclic PROTAC is considered (Case Study 1),^56^ when the degraded proteins differ only by three residues (Case Study 2),^54^ when direct protein-protein interactions are known to be weak (Case Study 4),^48^ when the PROTAC is known to hit multiple protein targets (Case Study 5),^62^ or when the PROTACs themselves show subnanomolar potency (Case Study 6).^64^ As was shown, accurate results can result even if the input binary structures themselves must be modeled through small molecule docking, even in the (unfortunately common) scenario where the crystallographic precedent is incomplete, such as for the pan-kinase binding moiety incorporated into the PROTAC in Case Study 5. For these inherently more difficult cases, useful accuracy perhaps requires a more thorough procedure to generate a diverse array of input structures, as might be expected.

There are, of course, some exceptions to the overall accurate modeling results discussed in this work. Case Study 3 highlighted one potential challenge for Method 4B: a simple population-based scoring metric, even utilizing the double clustering procedure of Method 4B, generally tends to favor a longer or more flexible PROTAC over a shorter or rigidified alternative, by a simple “more shots on goal” logic. However, particularly for early-phase PROTAC discovery efforts, this phenomenon is perhaps not at all artificial: that is, an unoptimized PROTAC may indeed be more likely to successfully bring two proteins together if a longer or more flexible linker provides more opportunities to achieve the requisite proximity. Moreover, the correlation between PROTAC conformational flexibility and final predicted degradation efficiency found in Case Study 3 is not ironclad – for example, Case Study 4 showed a diminution of this effect for particularly long PROTACs (Figure 4a), while Case Study 6 demonstrated that Method 4B can successfully discriminate between better and worse degraders even if the number of PROTAC conformations is similar (*cf*. PROTACs 29, 35, 36, 37, and 32 of Figure 6b, which all use ensembles of 25-35 PROTAC conformations). It seems that the additional machinery in Method 4B, particularly the protein-based filters that enforce a maximum allowed interfacial surface area of 670Å^2^, offer something of a safeguard against overscoring flexible PROTACs, as these filters are agnostic of the PROTAC entirely. Nonetheless, caution and good sense should certainly be exercised if modeled results predict that longer PROTACs are invariably better, with no predicted upper limit. Again we note the parallel to small molecule docking scores, which also correlate with molecular size.^61^

The second set of (somewhat) inaccurate predictions was revealed in Case Study 7, where degraders utilizing the E3 ligase cIAP1 were ranked less accurately than were the systems considered in other case studies. However, in a sense this finding is oddly reassuring, in that there is little reason to *a priori* expect that the criteria used to judge the acceptability of a modeled VHL- or cereblon-containing ternary complex should apply to complexes made from *any* arbitrary E3 ligase. Nonetheless, as *some* of the experimental trends were successfully recapitulated by the Method 4B modeling in Case Study 7, it may be that some similarities can be found across ternary complex structures containing VHL, cereblon, and cIAP1. For example, inspection of multiple cIAP1 crystal structures (Figure S3) reveals that the E3 ligase recruiting moiety binds to a pocket found on the *side* of the cIAP1 BIR3 domain, rather than to a pocket located at one *end* of the enzyme, as in VHL and cereblon (Figure S1). Thus, the requirement enforced by Method 4B for a direct protein-protein interfacial surface area of <670Å^2^ in a “crystal-like” ternary complex may be generalizable to other E3 ligases. Intriguingly, it can also be observed that the degraded kinases in Case Study 5 interact with cereblon via a kinase binding moiety bound in the canonical ATP binding site, which of course is located in a cleft between the N- and C-terminal lobes rather than at one “end” of the kinase. Thus, it may be that effective PROTAC-mediated degradation requires at least one of the two proteins in the three-body ternary complex to approach the other in an end-on fashion, which VHL and cereblon (Figure S1) naturally accommodate, but which can perhaps also be ensured by the approach orientation of the target protein. Clearly, this possibility requires further exploration, but it seems sensible from a simple packing perspective: in order for two pockets to be bridged by a PROTAC of reasonable size, they must be able to get close to one another, which would be facilitated most by an end-on approach of at least one of the proteins. Such a consideration would have far-reaching consequences for the choice of an E3 ligase to use to effect degradation, once the library of choices has expanded to afford the luxury of such a choice.^19^

Regardless of whether this hypothesis is correct, it is apparent from the development of (and results predicted by) Method 4B that the nature, quality, and diversity of the protein-protein docking poses are critical – the improved effectiveness of reproducing the structure of 6BOY once the more thorough “Multidock” procedure is invoked (Figure 3) is a clear indication of this fact. In a sense, the modeling procedure of Method 4B is somewhat roundabout: initially the method constructs large, diverse ensembles of both protein-protein docked poses and PROTAC conformations, but then various filters and clustering procedures are used to winnow these diverse ternary complex models down to a more focused description, which hopefully resembles the actual ternary complex geometry. In other words, variety is sought during the input phase, only to be shed when the final output is assembled.

However, this roundabout procedure is necessary for two reasons. The first is that the scores used to judge the quality of a predicted protein-protein interface, even including the physics-based score (forcefield) score used in MOE, are simply not accurate enough to identify the ephemeral interfaces in a ternary complex that never naturally occur without the intervention of a PROTAC. Indeed, the area of designing, judging, and improving protein-protein docking algorithms and scoring functions^65^ is largely based on the ability to reproduce *in silico* known structures of natively interacting proteins, which clearly differs from being able to correctly describe the *ad hoc* interface that is created in a PROTAC-mediated ternary complex. Thus, Method 4B favors adding and then removing diversity simply because the alternative – to naively take some slice of “high” scoring protein-protein docked poses – is almost certain to disappoint. Moreover, it is important to note that the input structures provided to protein-protein docking in a PROTAC-mediated scenario (*i*.*e*., the two binary target+binder and ligase+binder complexes) generally incorporate the *unbound* form of each protein, which is a well-known challenge for protein-protein docking algorithms.^66^

Secondly, Methods 4, 4B, and 5 were designed to ultimately judge the *compatibility* between a given PROTAC and the two input protein complexes, as shown by, for example, the use of Core RMSD as a filtering criterion. Even if a scoring function could be developed to accurately score the unoptimized and generally nonspecific interface between a target protein and an E3 ligase, the poses scored most favorably would almost certainly change once the PROTAC is introduced to the system. Indeed, one of the key advantages (and challenges!) of a PROTAC-mediated TPD approach is that even slight changes in the PROTAC can lead to decidedly different target-ligase orientations, which in turn often enables a degree of degradation selectivity, even amongst closely related targets, as has been demonstrated for both VHL-^67^ and cereblon-mediated degradation.^55^ If an input ensemble of protein-protein docked poses were *a priori* limited based on a protein-protein docking score, even an incredibly accurate one, the ability to identify novel poses only favored once a PROTAC with a particular conformational ensemble is present would be severely curtailed. Importantly, we designed Methods 4, 4B, and 5 to accept input protein-protein docked ensembles generated by essentially any software, so specific techniques that prove especially well-suited to the unique challenges of predicting PROTAC-mediated ternary complexes can be identified.

Finally, it should be noted that all of the Methods we have developed, both herein and in our earlier work,^41^ are of use to *screen* PROTACs. To be precise, the *user* must provide as input one or more prospective PROTACs to analyze. To facilitate such efforts, these tools were designed to be both modular and embarrassingly parallel: for example, the conformational ensemble of the PROTAC TL12-186 considered in Case Study 5 was generated *once*, and then copies of this ensemble were passed to simultaneously running simulations exploring different kinases and different target binding pose orientations. Similarly, the ensemble of BTK+binder docked against cereblon+binder in Case Study 4 was generated once, and then copies of this protein-protein docked ensemble were used in multiple independent simulations, to predict the relative degradation efficacy of PROTACs 2-12 all at once. However, none of our Methods will attempt to generate any *new* PROTACs *de novo* – in other words, these tools do not propose or *design* novel PROTACs suitable for a given system. Nonetheless, the modeling results provided by our methods, particularly the double clustered ensembles produced by Method 4B, can be used as starting points for traditional structure-based iterative design, as has already been successfully accomplished starting from a crystallographic ternary complex.^52,54–56^ The advantage of using a computationally modeled starting point instead, of course, is greater expediency and diminished cost.

## ASSOCIATED CONTENT

The following files are available free of charge.

Additional development details for Method 4B and Method 5. Method 5 Results for Case Studies 2 and 3. Data and supplementary results for Case Study 4. Supplementary data for Case Study 7. (PDF)

## ACKNOWLEDGMENT

The authors acknowledge many helpful discussions, suggestions, and feedback from: Emel Ficici, Yilin Meng, and Ye Che (Pfizer), Veer Shanmugasundaram (formerly Pfizer/currently Celgene), Theresa Johnson and Kathleen Dreher (EMD-Serono), Michelle Lamb and Scott Pedersen (AstraZeneca), Cen Gao (formerly Eli Lilly), Kiyo Omoto and Nick Calandra (H3 Biomedicine), Yelena Arnautova (X-Chem), Sarah Meitz and Philipp Schmalhorst (Boehringer Ingelheim), and Li Xiao (Merck).

## ABBREVIATIONS

PROTAC: proteolysis-targeting chimera
TPD: targeted protein degradation
CDK: cyclin-dependent kinase
ATP: adenosine triphosphate
VHL: von-Hippel Lindau
PK: pharmacokinetics
PD: pharmacodynamics
PDB: Protein Data Bank
RMSD: root-mean-square deviation
MOE: Molecular Operating Environment
SVL: Scientific Vector Language
CAPRI: Critical Assessment of Predicted Interactions
PEG: polyethylene glycol
TR-FRET: time-resolved fluorescence energy transfer
BET: bromo- and extra-terminal
BD: bromodomain
IAP: inhibitor of apoptosis protein
SNIPER: specific and nongenetic inhibitor of apoptosis protein-dependent protein eraser
AR: androgen receptor

