## Supporting Information for "Improved Accuracy for Modeling PROTAC-Mediated Ternary Complex Formation and Targeted Protein Degradation via New *In Silico* Methodologies"

### Additional Development Details of Method 4B and Method 5

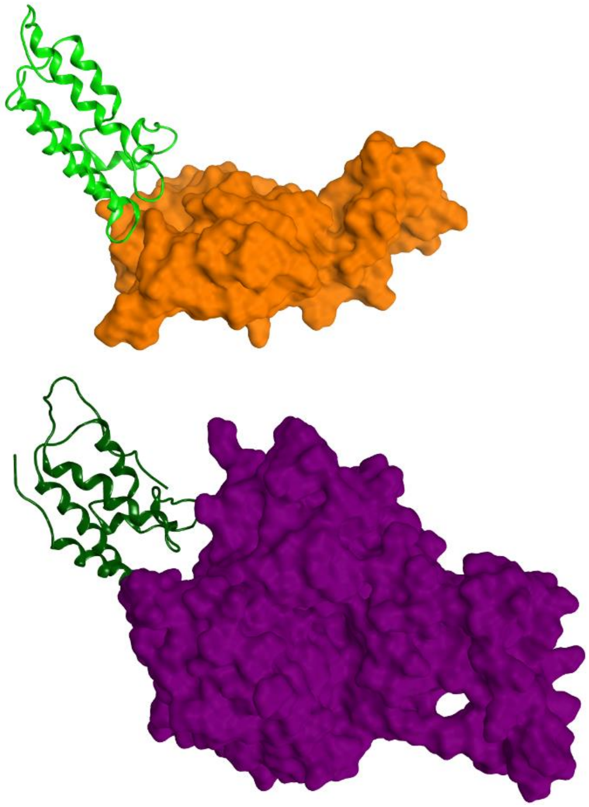

**Figure S1.** End-on target binding arrangements illustrated in ternary complex crystal structures of (top, 6HAX) SMARCA2$^{BD}$ (green helices) with VHL (orange surface) and (bottom, 6BN7) Brd4$^{BD1}$ (dark green helices) with cereblon (purple surface).

**Table S1. Hit Rates for the Validation Set of Known Ternary Complex Structures of Table 1 (Method 4B), with and without Clustering, as Measured by Two Criteria**

| | Crystal-Like/Total (Hit Rate) | | High/Medium/Acceptable (Hit Rate)[a] | |
|---|---|---|---|---|
| PDB | Unclustered | Clustered | Unclustered | Clustered |
| 5T35 | 1047/1692 (61.9%) | 979/1040 (94.1%) | 50/950/198 (70.8%) | 49/886/98 (99.3%) |
| 6BN7 | 15/390 (3.8%) | 11/101 (10.9%) | 0/5/15 (5.1%) | 0/3/11 (13.9%) |
| 6BOY | 4/1063 (0.4%) | 0/87 (0.0%) | 0/4/0 (0.4%) | 0/0/0 (0.0%) |
| 6BOY MD[b] | 229/1604 (14.3%) | 171/691 (24.7%) | 0/17/157 (10.8%) | 0/8/126 (19.4%) |
| 6HAX | 143/526 (27.2%) | 143/464 (30.8%) | 88/36/71 (37.1%) | 88/27/71 (40.1%) |
| 6HAY | 79/420 (18.8%) | 75/313 (24.0%) | 0/36/144 (42.9%) | 0/36/144 (57.5%) |
| 6HR2 | 65/447 (14.5%) | 49/172 (28.5%) | 0/67/32 (22.1%) | 0/67/5 (41.9%) |

[a]Across all three categories [b]Using the “multidock” procedure (see main text)

### Method 5 Results for Case Studies 2 and 3 (Gadd *et al.*[1] and Zengerle *et al.*[2])

**Table S2. Predictions for Wild Type and QVK Triple-mutant of Brd4$^{BD2}$ using Method 5 (Case Study 2)**

| | RMSD_RF | ligand_E | RMSD_RF_PP | ligand_E_PP |
|---|---|---|---|---|
| Wild Type | 4 | 43 | 4 | 20 |
| QVK | 1 | 32 | 0 | 21 |

**Table S3. Predictions for Three PROTACs and Three Brd Targets using Method 5 (Case Study 3)**

| | RMSD_RF | | | RMSD_RF_PP | | |
|---|---|---|---|---|---|---|
| | MZ1 | MZ2 | MZ3 | MZ1 | MZ2 | MZ3 |
| Brd4$^{BD2}$ | 2 | 1 | 0 | 2 | 1 | 0 |
| Brd3$^{BD2}$ | 7 | 5 | 5 | 7 | 5 | 4 |
| Brd2$^{BD2}$ | 3 | 1 | 2 | 3 | 1 | 2 |

| | ligand_E | | | ligand_E_PP | | |
|---|---|---|---|---|---|---|
| | MZ1 | MZ2 | MZ3 | MZ1 | MZ2 | MZ3 |
| Brd4$^{BD2}$ | 43 | 47 | 40 | 21 | 18 | 18 |
| Brd3$^{BD2}$ | 50 | 50 | 47 | 36 | 34 | 31 |
| Brd2$^{BD2}$ | 36 | 45 | 39 | 22 | 24 | 20 |

### Data and Supplementary Results for Case Study 4 (Huang *et al.*[3])

**Table S4. Input Crystal Structures and their Ligands selected for the Kinases of Case Study 4**

| Kinase | PDB | Ligand |
|---|---|---|
| AAK1 | 5TE0 | 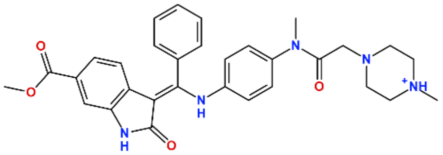 |
| AURKA | 3E5A | 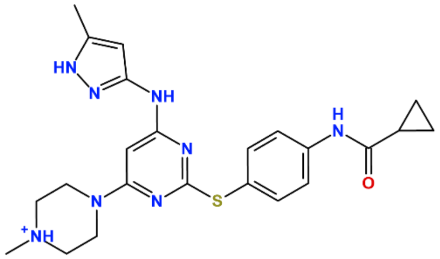 |
| AURKB | 5K3Y | 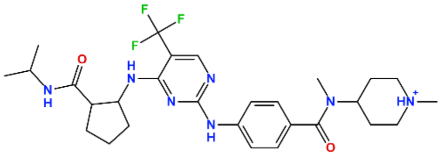 |
| BTK | 4OT5 | 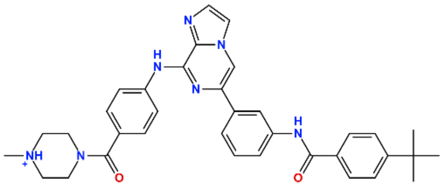 |
| CDK2 | 2C4G | 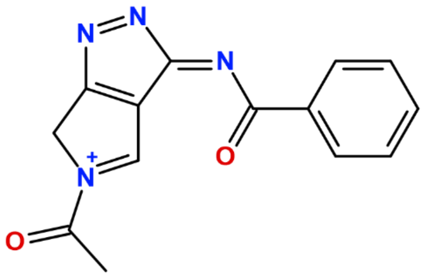 |
| CDK5 | 1UNL | 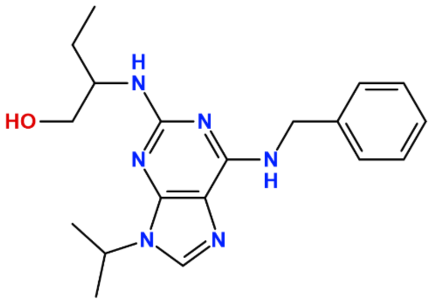 |

| Kinase | PDB | Ligand |
|---|---|---|
| CDK6 | 2EUF | 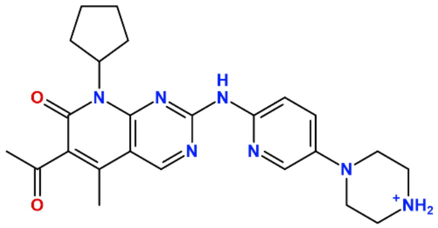 |
| CDK7 | 1UA2 | 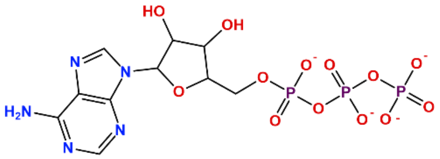 |
| CDK9 | 3BLQ | 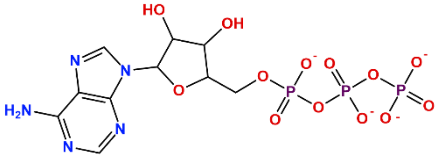 |
| CDK12 | 6B3E | 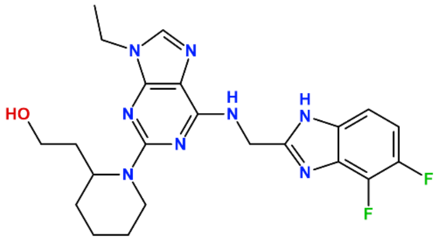 |
| FES | 4E93 | 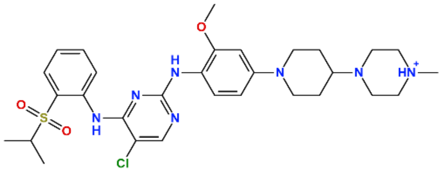 |
| FLT3 | 5X02 | 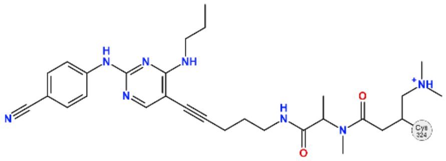 |
| GAK | 4C57 | 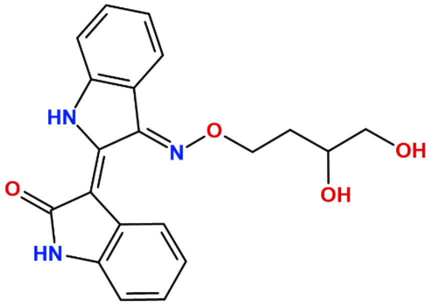 |

| Kinase | PDB | Ligand |
|---|---|---|
| INSR | 3EKN | 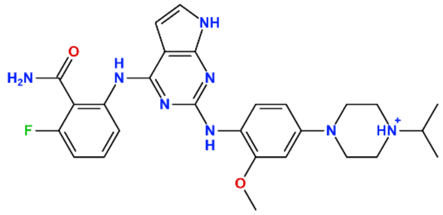 |
| ITK | 4L7S | 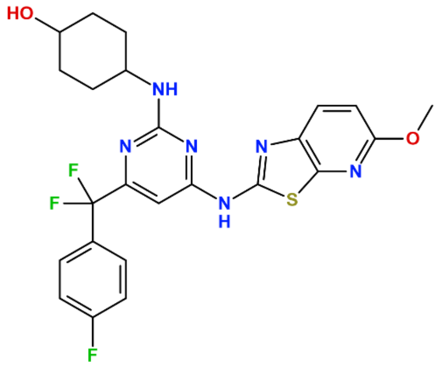 |
| JAK1 | 5E1E | 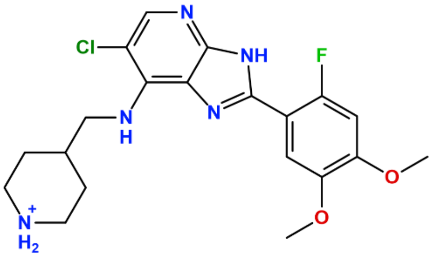 |
| MARK2 | 5KZ7 | 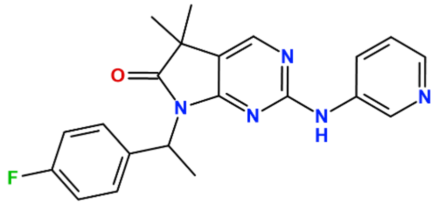 |
| PRKAA1 | 5T5T | 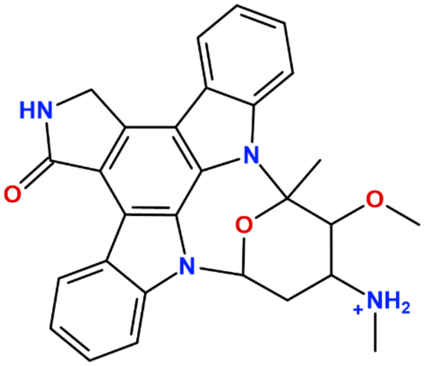 |

| Kinase | PDB | Ligand |
|---|---|---|
| PTK2 | 4GU6 | 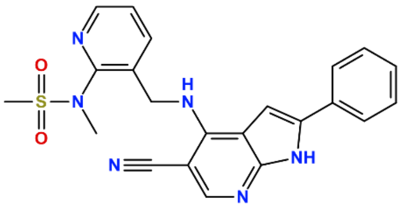 |
| PTK2B | 3ET7 | 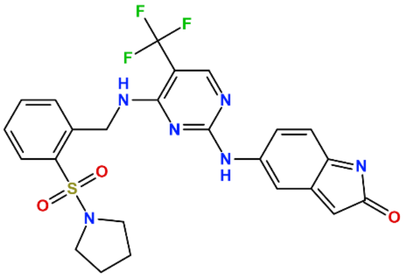 |
| SLK | 2J7T | 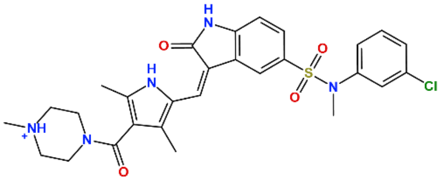 |
| TAK1 | 4L3P | 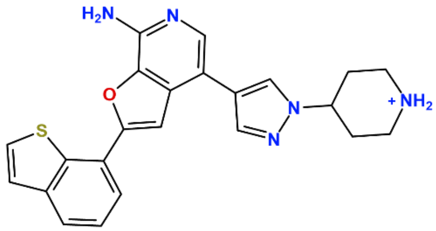 |
| TBK1 | 4IWQ | 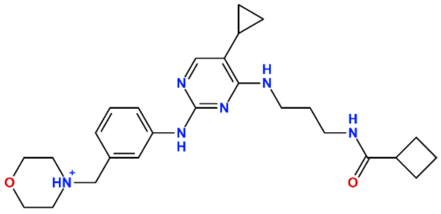 |
| ULK1 | 4WNP | 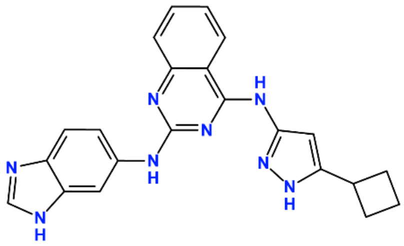 |
| WEE1 | 5VC3 | 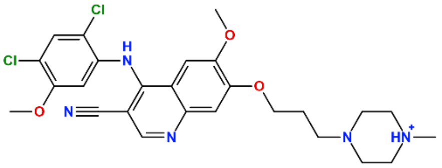 |

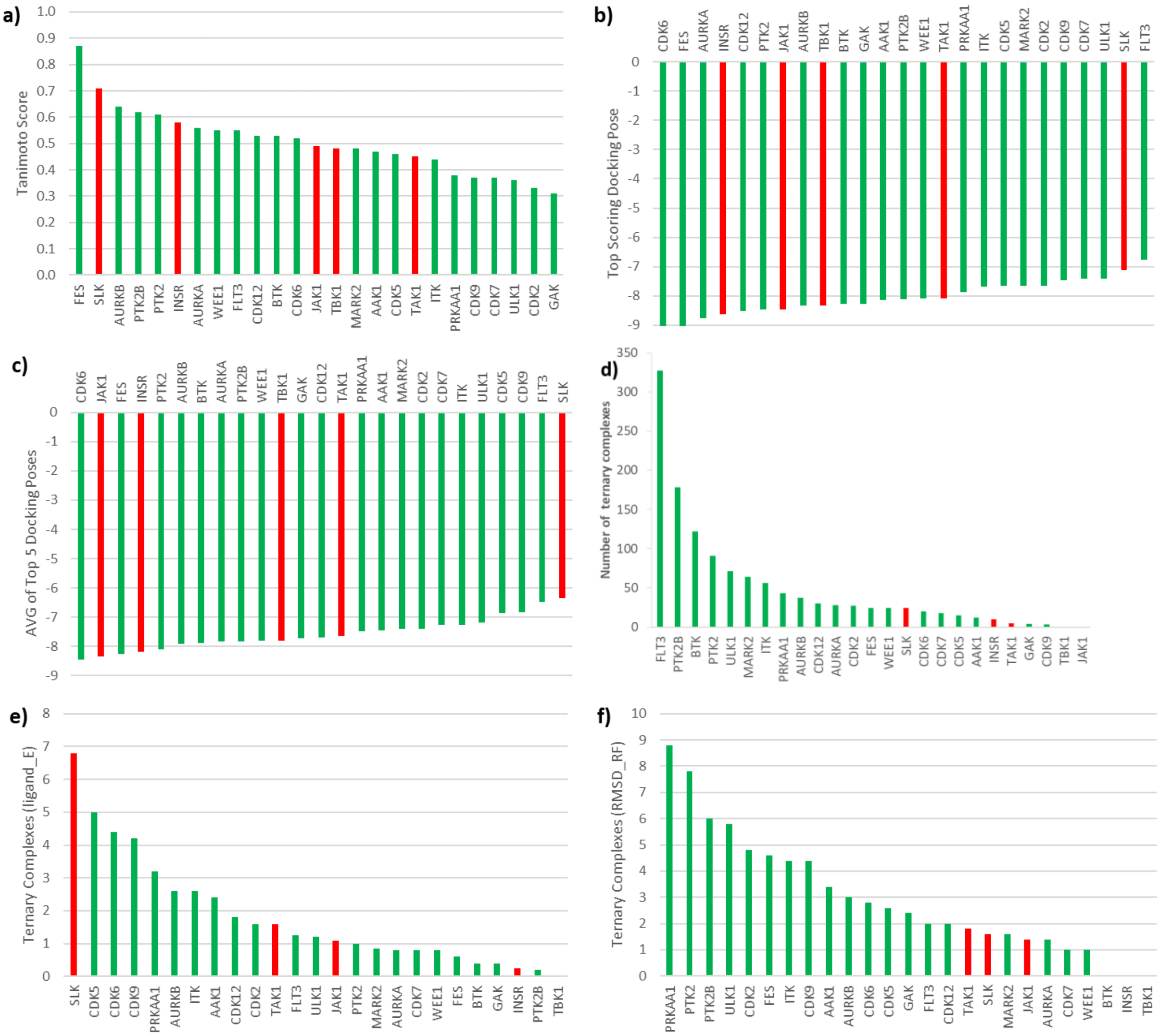


**Figure S2.** a) Tanimoto similarities between the kinase ligands of Table S4 and the pan-kinase binding moiety TL13-87 incorporated into the PROTAC TL12-186. Green bars indicate kinases that are degraded by TL12-186, while red bars show kinases that bind to the PROTAC but do not degrade. This same green/red coloring convention is used in b)-f) as well. b) Top GBVI/WSA dG score after TL13-87 is docked into the corresponding kinase pocket. c) Average of the scores for the top five manually identified docking poses for the corresponding kinases. d) Results from Method 4$^{D}$ (see Ref. 4) for the kinases of this study, illustrating how the requirement in Method 4$^{D}$ for direct target-E3 ligase hydrophobic-hydrophobic patch contact disfavors SLK (*cf*. Figure 5 in the main text of this work). e) Results with Method 5, using the ligand_E metric, averaged across the top five poses for each kinase. f) Results with Method 5, using the RMSD_RF metric, averaged across the top five poses for each kinase.

### Supplementary Data for Case Study 7 (Shibata *et al.*[5])

**Table S5. SNIPERs Modeled in this Work and their Experimental AR Expressions (% of Control)**

| SNIPER | Structure | Expt. AR Expression (% of Control) | Source of Expt. Value from Ref. 5 |
|---|---|---|---|
| 2 | 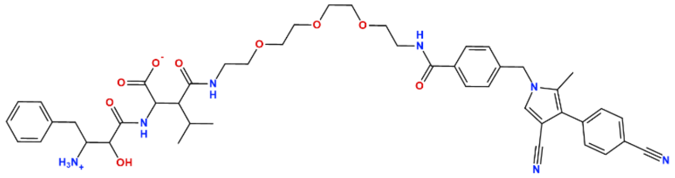 | 37<br>96 | Fig. 1C (30 µM)<br>Fig. 2A (3 µM)[a] |
| 23 | 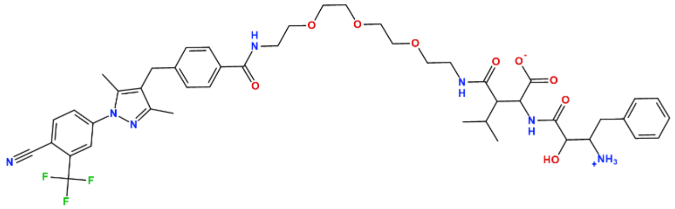 | 26 | Fig. 1C (30 µM) |
| 26 | 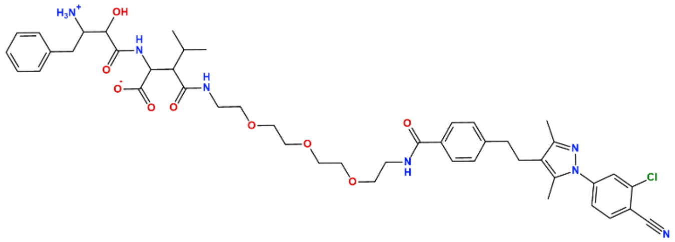 | 31 | Fig. 1C (30 µM) |
| 8 | 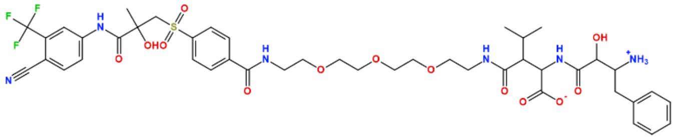 | 67 | Fig. 1C (30 µM) |
| 27 | 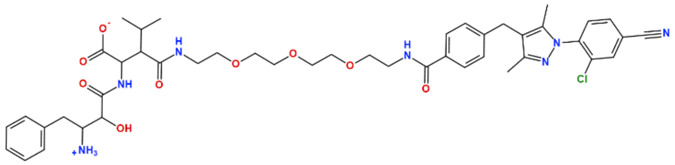 | 88 | Fig. 1C (30 µM) |
| 31 | 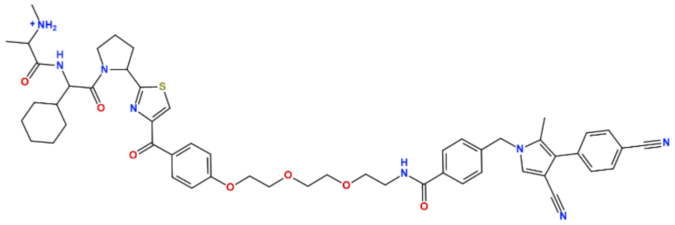 | 30 | Fig. 2B (3 µM) |
| 60 | 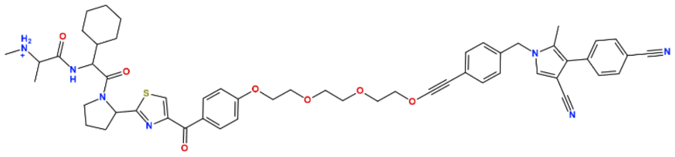 | 24 | Fig. 2B (3 µM) |

| SNIPER | Structure | Expt. AR Expression (% of Control) | Source of Expt. Value from Ref. [5] |
|---|---|---|---|
| 70 | 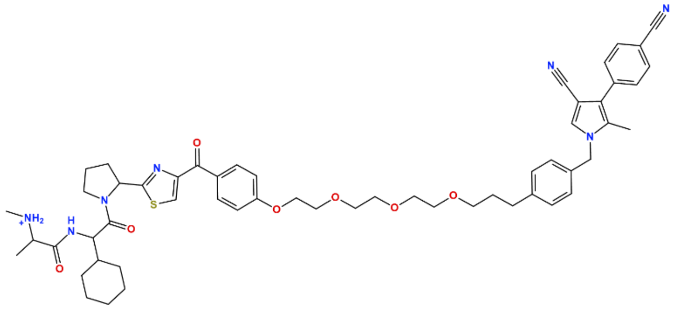 | 18 | Fig. 2B (3 μM) |
| 53 | 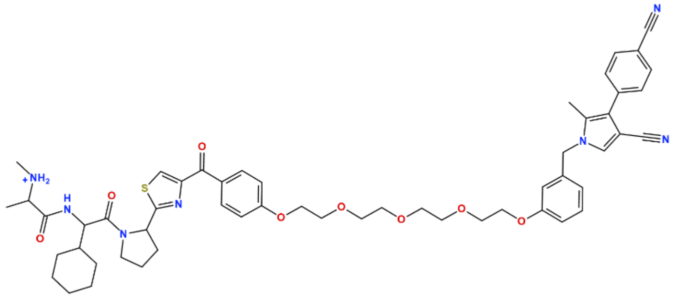 | 36 | Fig. 2C (3 μM) |
| 65 | 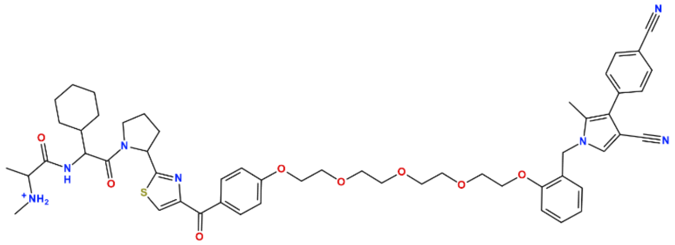 | 80 | Fig. 2C (3 μM) |
| 52 | 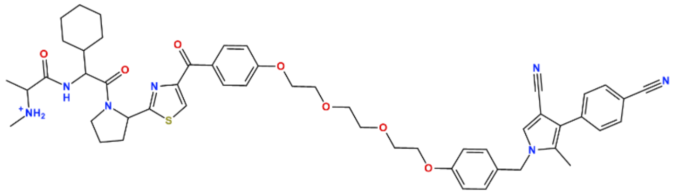 | 12 | Fig. 3A (3 μM) |
| 171 | 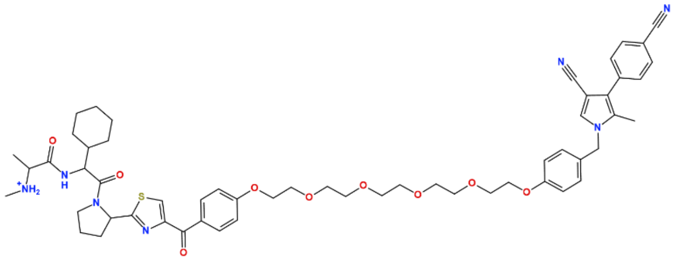 | 9 | Fig. 3A (3 μM) |
| 51 | 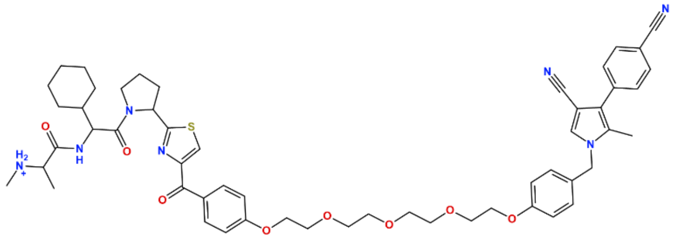 | 23 | Fig. 3B (3 μM) |

| SNIPER | Structure | Expt. AR Expression (% of Control) | Source of Expt. Value from Ref. [5] |
|---|---|---|---|
| 66 | 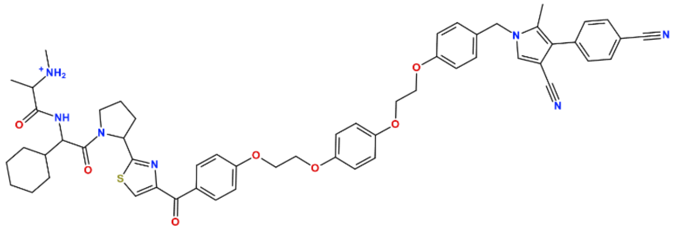 | 107 | Fig. 3B (3 μM) |
| 67 | 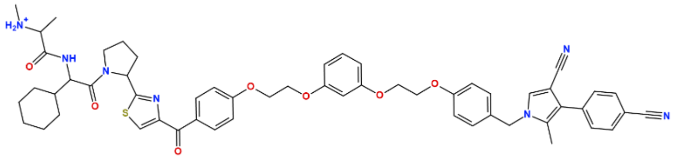 | 90 | Fig. 3B (3 μM) |
| 68 | 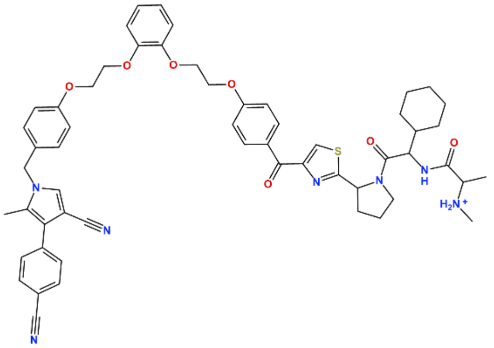 | 101 | Fig. 3B (3 μM) |

[a]For comparison with SNIPER-31 in Figure 7a in the main text

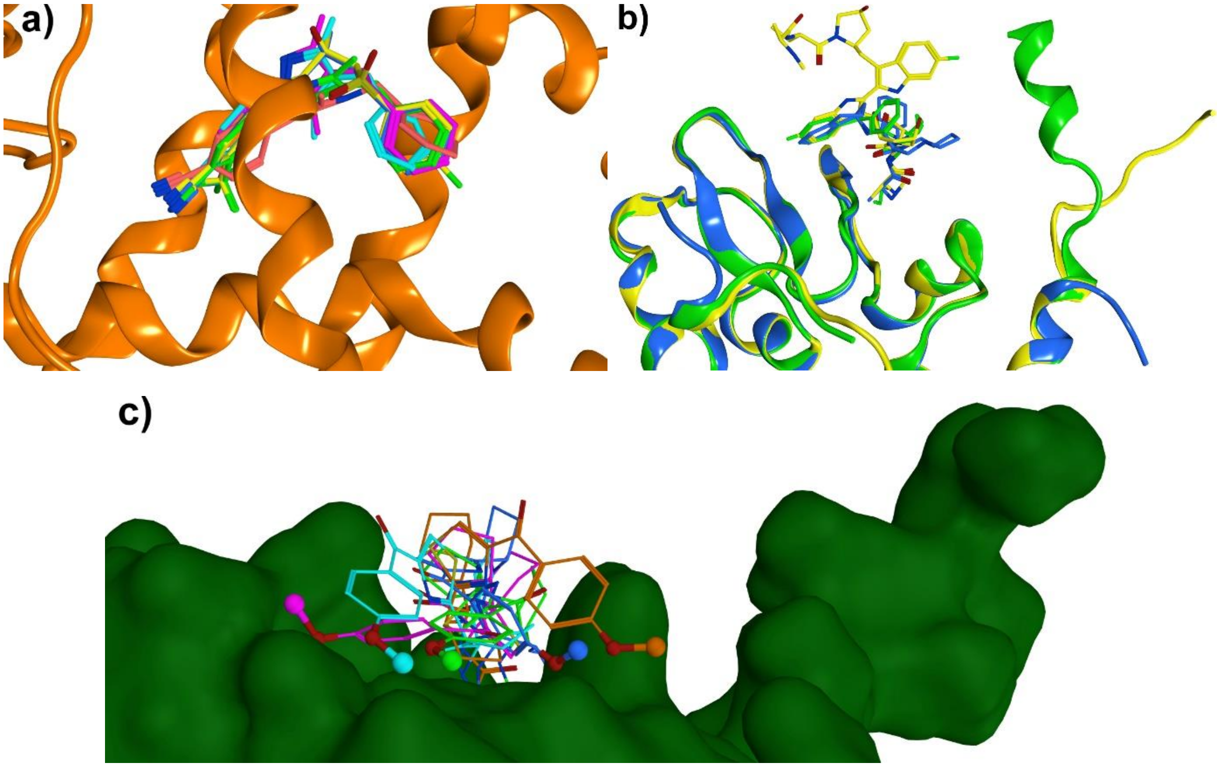


**Figure S3.** a) Common orientation of the ligands in the binding pocket of the androgen receptor (PDB ID: 1Z95) from the crystal structure (yellow) and for docked poses of the target binding moieties of SNIPER-2 (salmon), SNIPER-8 (blue), SNIPER-23 (green), SNIPER-26 (magenta), and SNIPER-27 (cyan). b) Superposed crystal structures of the cIAP1-BIR3 domain used to generate input poses for the E3 ligase+binder complex: 3MUP (green), 4HY4 (blue), and 4KMN (yellow). The variability in the conformations for the C-terminii are evident on the right. C) Five poses of Compound 24 (see text) docked into the ligand binding site of cIAP1 from 4KMN. The methoxy where the SNIPER linkers attach are indicated with small spheres.

### References used in the Supporting Information
